# A Unique Role for the Hippocampus in Cue Integration During Human Spatial Navigation

**DOI:** 10.1101/2025.05.14.654008

**Authors:** Ziwei Wei, Thomas Wolbers, Xiaoli Chen

## Abstract

It is a central question in cognitive neuroscience how the brain integrates diverse sensory inputs to improve cognitive performance. This study investigated this question in the domain of human spatial navigation using high-field fMRI, a novel navigation task, and desktop virtual reality. Participants learned and retrieved spatial locations using landmarks alone (landmark condition), visual self-motion cues alone (i.e., optic flow; self-motion condition), or both cues together (combination condition). Behaviorally, participants benefited from cue integration. fMRI analyses revealed a cue integration effect in the hippocampus, which displayed positional coding only in the combination condition, and only in participants who showed behavioral evidence of cue integration. Notably, the posterior hippocampus exhibited stronger positional coding in the combination condition than in both single-cue conditions. Additionally, positional coding in the hippocampus predominantly reflected actual, rather than self-reported, locations, suggesting relatively early involvement in the cue integration process. These findings suggest that the hippocampus supports precise memory-guided navigation by integrating different spatial cues, a function beyond simple spatial representation. More broadly, this work helps reconcile the longstanding debate regarding the hippocampus’s role in spatial navigation versus episodic and relational memory, facilitating a unified understanding of hippocampal function.

## Introduction

Effective behavior in complex environments depends on the brain’s ability to combine multiple, often noisy, sources of information. Cue integration is a fundamental cognitive process that supports perception, decision-making, and memory by reducing uncertainty and enhancing reliability across sensory and informational domains. In spatial navigation, accuratly localizating critical positions in the environment, such as food sources or home bases, is a key function of the cognitive map – a mental representation of space that supports adaptive navigation. Navigators often face a multitude of spatial cues, each contaminated by sensory noise. This noise can be reduced through the integration of different cues (Cheng, et al., 2007), leading to a more robust cognitive map and more accurate navigation.

Behavioral studies have demonstrated that people can integrate different spatial cues (e.g., landmarks and self-motion cues) to improve spatial localization performance in navigation tasks (for a review, see Newman et al., 2023). However, how the brain accomplishes this integration remains poorly understood. In humans, Huffman and Ekstrom (2019) investigated neural representations of locations by varying the availability of body-based self-motion cues in the presence of visual information. Their findings provide evidence for cue-independent spatial representations across a large brain network, including the retrosplenial cortex (RSC) and the hippocampus. Recently, we have characterized neural representations of spatial locations encoded from landmark and visual self-motion cues, detecting both cue-specific and cue-independent spatial representations in the entorhinal cortex (EC) (Chen et al., 2019) and RSC (Chen et al., 2024, 2025). Despite these advances, no human studies have yet investigated the neural basis of spatial cue integration during navigation.

In contrast, the interaction between different spatial cues during navigation has been intensively studied in non-human animals. Two lines of such research are related to spatial cue integration. First, neural responses of location-sensitive cells, such as hippocampal place cells and entorhinal grid cells, often show a compromise between conflicting spatial cues, with firing fields positioned between the locations defined by the conflicting cues (Campbell et al., 2021; Chen et al., 2013; Gothard et al., 1996; Jayakumar et al., 2019; Madhav et al., 2024). However, this compromise may reflect a mechanism different from cue integration. For example, in cue alternation, navigators rely exclusively on one cue for a subset of trials and the other for the remainder, whereas in cue integration, navigators combine different cues within individual trials (Nardini et al., 2008). Like cue integration, cue alternation leads to compromises between different cues. However, unlike cue integration, cue alternation does not enhance localization precision. Second, adding an additional spatial cue often improves the quality of positional coding in place cells (Quirk et al., 1990) and grid cells (Chen et al., 2016), for example, by reducing the firing field size and improving firing stability across sessions. However, these improvements do not necessarily indicate cue integration, as these studies did not compare the double-cue condition with all relevant single-cue conditions (Chen et al., 2017). Therefore, the observed improvements in positional coding may simply reflect exclusive reliance on the newly added, more precise cue. In sum, while electrophysiological studies have provided valuable insights into the neural mechanisms underlying spatial cue interaction, they do not offer direct evidence of how the brain integrates spatial cues during navigation.

In this study, we examined how the brain integrates landmarks and self-motion cues in during navigation. Landmarks and self-motion cues are two fundamental sources of spatial information, recruiting relatively independent neural and cognitive processes (Chen et al., 2019; Etienne et al., 1996). Humans can combine these cues to enhance navigation performance (Chen et al., 2017). Young healthy participants encoded and retrieved four fixed locations along a linear track, using either landmarks alone, visual self-motion cues alone (i.e., optic flow), or a combination of both in a desktop virtual reality environment while undergoing 3T fMRI scanning. Focusing on the medial temporal lobe (MTL) and retrosplenial cortex (RSC) – regions of interest in our previous investigations of single-cue spatial navigation (Chen et al., 2019, 2024, 2025) – we detected neural activity reflecting cue integration exclusively in the hippocampus. This finding underscores the hippocampus’s unique contributions to spatial cue integration during navigation.

**Figure 1.**
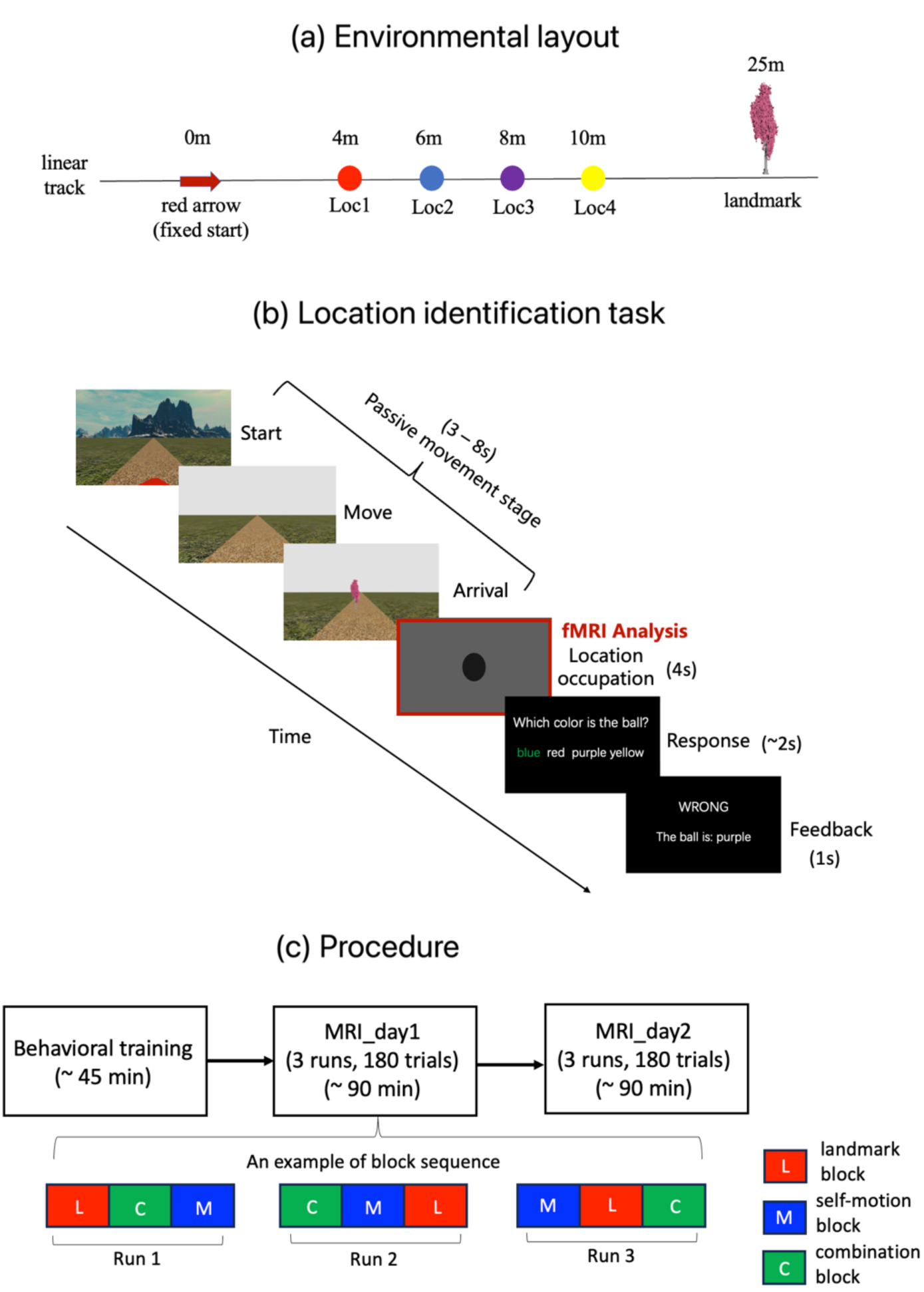
Environmental layout, experimental task, and procedure. **(a)** The environmental layout, including an arrow, a tree, and four differently colored balls on poles positioned at test locations Loc1-Loc4 along a linear track. **(b)** Location identification task time course, consisting of four stages. In the passive movement stage, participants started at the arrow (0m), and were passively transported to one of the four test locations. The tree appeared at the end of the movement (depicted larger for illustration). In the location occupation stage, participants’ first-person perspective was smoothly tilted downward and fixed on the ground for 4 seconds. In the response stage, participants recalled the color of the ball positioned at the location within 20 seconds. In the feedback stage, feedback was given, along with the correct answer if incorrect. The balls remained invisible throughout the trial. The background environment disappeared once the movement started. All fMRI analyses focused on the 4-second location occupation stage, where visual inputs were identical across cue conditions. **(c)** Procedure. On Day 1 (Pre-scan), participants were trained on the location identification task. On Days 2 and 3 (MRI_day1 & MRI_day2), they performed the location identification task during scanning in a 3T MRI scanner.

## Results

Thirty-five healthy young adults (20 males, mean age = 23.8 years) participated in this study. They first learned the positions of four locations arranged along a linear track in a desktop virtual-reality environment (Figure 1a). Next, they performed a location identification task (Figure 1b) during 3T MRI scanning, which was conducted over two sessions on two separate days. In each trial of this task, they were passively moved to one of the four test locations, stayed there for 4 seconds, and then identified the location. This task included three cue conditions: landmark (a tree), self-motion, and combination. To match the sensory inputs across conditions as much as possible, both the landmark and self-motion cues were provided in all three conditions. In the two single-cue conditions, the landmark was adjusted to create spatial conflicts between self-motion and landmark cues, meaning the correct target location defined by the two cues differed (Methods). Participants were instructed to rely exclusively on either the landmark cue (landmark condition) or self-motion cues (self-motion condition). Feedback was provided according to the task-relevant cue type in the condition to reinforce the designated navigation strategy. In the combination condition, the landmark was positioned at its original position, allowing participants to utilize both cue types to identify the test location. Feedback was also provided in the combination condition.

### Cue Integration Enhances Behavioral Accuracy in Location Identification

Our primary focus was whether participants exhibited cue integration by showing better performance in the combination condition compared to each single-cue condition. Behavioral accuracy was submitted to repeated measures ANOVA with cue type, test location and scanning day as independent variables (Figure 2a). The main effect of cue condition was significant (F(2, 68) = 67.100, p < 0.001, 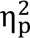 = 0.664, BF_10_ = 6.314). Planned comparisons showed higher behavioral accuracy in the combination condition compared to both the landmark condition (t(34) = 4.303, p < 0.001) and the self-motion condition (t(34) = 11.808, p < 0.001). A closer look at individual test locations showed that the simple effect of cue condition was significant at all four locations (ps < 0.001) (Figure 2a, right panel). The pattern of results was consistent across test locations, with the combination condition outperforming both single-cue conditions (ts > 3, ps < 0.015), even though the difference between the combination and landmark conditions did not reach statistical significance at Loc3 (t(34) = 1.911, p = 0.194) and Loc4 (t(34) = 1.337, p = 0.571).

We matched sensory inputs between single-cue conditions and the combination condition by presenting both cue types in the single-cue conditions, where participants were instructed to rely on one cue type (task-relevant) and ignore the other one (task-irrelevant). If participants adhered to this instruction, their behavioral accuracy should decrease as the test location gets farther away from the respective anchoring points of landmark-based navigation (i.e., the landmark) (Chamizo et al., 2006) and path integration (i.e., the arrow) (Loomis et al., 1999). The two anchoring points were positioned at the two ends of the linear track, predicting an interaction between cue type and the linear trend of the test location along the track. This interaction was significant (t(34) = 9.398, p < 0.001). Behavioral accuracy decreased linearly as locations became farther away from the tree in the landmark condition (t(34) = 2.487, p = 0.013), and from the arrow in the self-motion condition in the self-motion condition (t(34) = 8.799, p < 0.001). However, one deviation emerged: accuracy at Loc1 was higher than at Loc2 in the landmark condition (t(34) = 2.751, p = 0.006), contrary to the expected trend. This result likely reflects reduced margin for error at this location, given that it had only one adjacent location (i.e., Loc2). Supporting this explanation, cognitive modeling showed that positional precision followed expected cue-driven spatial patterns (Figure 2b, right panel; interaction: t(102) = 8.882, p < 0.001). Moreover, cue integration was evident even at Loc1, where accuracy in the combination condition significantly exceeded both single-cue conditions (ps < 0.001). These findings are consistent our previous report where only the task-relevant cue was available in single-cue conditions (Chen et al., 2024), indicating that participants understood and followed experiental instructions.

In summary, participants displayed improved performance in the combination condition compared to both single-cue conditions, demonstrating integration of landmarks and self-motion cues. This cue integration effect is validated by participants’ behavior in single-cue conditions, which aligned with the expected navigation strategies for landmark-based navigation and path integration, thus confirming their suitability as baselines for assessing cue integration effect.

**Figure 2.**
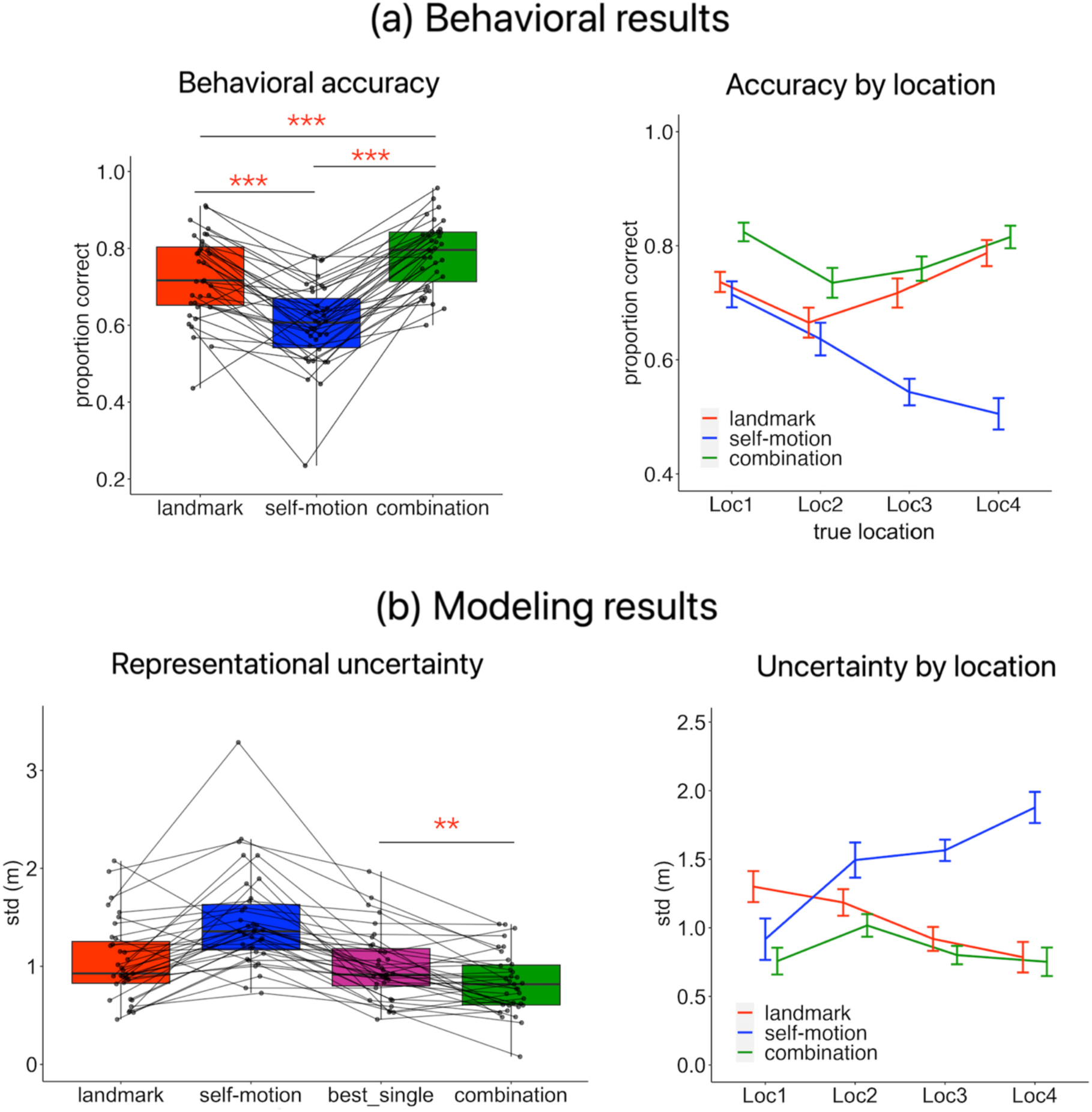
Behavioral and modeling results. **(a)** Behavioral results. Left panel displays boxplots of accuracy by cue condition. Black dots/lines represent individual participants. Right panel shows accuracy by cue type and test location. **(b)** Modeling results. Left panel shows representational uncertainty (averaged across test locations) by cue condition, along with the “best_single” representational uncertainty (lowest uncertainty of single-cue conditions). Right panel displays representational uncertainty across test locations. “**” indicates p_1-tailed_ < 0.01; “***” indicates p_1-tailed_ < 0.001.

### Cue Integration Enhances Precision of Positional Representations

While behavioral results showed improved performance in the combination condition, we next asked whether this enhancement was due to reduced representational uncertainty. To address this question, we fit participants’ responses with an extended signal-detection model, which disentangles representational uncertainty from response criterion and attentional lapse (Methods – Cognitive modeling: Extended signal detection theory).

As shown in Figure 2b (left panel), mean representation uncertainty was significantly lower in the combination condition compared to the landmark condition (t(34) = -4.023, p < 0.001, BF_10_ = 93), self-motion condition (t(34) = -8.609, p < 0.001, BF_10_ = 2.346*10^7^), and the single-cue condition with the lower uncertainty (vs. “best_single”, t(34) = -3.497, p = 0.001, BF_10_ = 24.622) (Scheller & Nardini, 2023). These findings indicate that participants integrated landmarks and self-motion cues to improve representational precision.

### Hippocampus Displays Positional Coding Reflecting Cue Integration

For fMRI analyses, we focused on BOLD signals from the location occupation phase of the location identification task, when participants remained stationary at the test location for four seconds (Figure 1b). Regions of interest (ROIs) included the retrosplenial cortex (RSC) and regions within the medial temporal lobe (MTL): the hippocampus, parahippocampal cortex (PHC), entorhinal cortex (EC), and perirhinal cortex (PRC) (Figure 3a). These areas are well-established for their roles in navigation and memory, and were also ROIs in our previous studies examining spatial representations derived from individual cues types (Chen et al., 2019, 2024, 2025). This consistent focus is pivotal for enhancing our understanding of the neural basis of spatial cue integration, which is likely distinct from single-cue navigation mechanisms. Crucially, all ROIs— except PRC—exhibited a successful navigation effect, with stronger activation in correct trials than incorrect ones (Figure S1). These results indicate that overall MTL and RSC were involved in the current task (Pessoa et al., 2002), validating our focus on these regions.

To identify brain regions involved in cue integration, we adopted the fMRI adaptation approach (Barron et al., 2016), which can assess neural coding of spatial relationships among the test locations: brain activation decreases as the spatial distance of successively visited test locations decreases (Chen et al., 2019, 2024; Morgan et al., 2011) (Methods). This approach allowed us to detect cue-integration-related neural activity, as improved precision of positional representations from cue integration should enhance the detectability of spatial coding in the brain.

A significant adaptation effect was detected exclusively in the hippocampus in the combination condition (t(34) = 2.915, p_1-tailed_ = 0.003, Cohen’s d = 0.493; Figure 3b, left panel), and the Bayes factor indicates strong evidence for the alternative hypothesis (BF_10_ = 12.722 > 10) (Jeffreys, 1961). Notably, this hippocampal effect remained significant after correcting for multiple comparisons across all 15 tests (p_corrected_ = 0.048; Methods). In contrast, the hippocampus displayed no significant adaptation effects in either the landmark condition (t(34) = 1.479, p_1-tailed_ = 0.074, Cohen’s d = 0.250, BF_10_ = 0.900) or the self-motion condition (t(34) = -0.852, p_1-tailed_ = 0.800, Cohen’s d = -0.292, BF_10_ = 0.106), and Bayes factors indicate evidence favoring the null hypothesis (BF_s10_ < 1). As visualized in Figure 2b (right panel), hippocampal activation in the combination condition increased linearly with inter-location distance, a pattern absent in the single-cue conditions.

Other ROIs did not show any significant adaptation effects in any cue conditions (ts < 1.15, ps1-tailed > 0.14, BF_s10_ < 0.55; Figure 3b, left panel). Given a previous study suggesting that alEC might be involved in cue integration (Doan et al., 2019), we conducted separate analyses for alEC and pmEC, as well as for each hemisphere. No significant adaptation effects were detected in any entorhinal subregions across cue conditions (ts < 1.37, ps > 0.18, BF_10_ < 0.43; but see Figure S2 & S3).

In sum, only the hippocampus displayed positional coding related to cue integration: this region displayed strong and robust location-based adaptation only in the combination condition but not in single-cue conditions. This hippocampal effect was not confounded by fMRI signal quality, as temporal signal-to-noise ratio was actually greater in RSC, PHC, and PRC than in the hippocampus, and also did not differ among cue conditions (Figure S4). These findings indicate that the hippocampus plays a unique role in spatial cue integration during navigation.

Overall, reliable adaptation-based spatial representations emerged in the hippocampus only when spatial cues were congruent, indicating the involvement of this region in cue integration.

**Figure 3.**
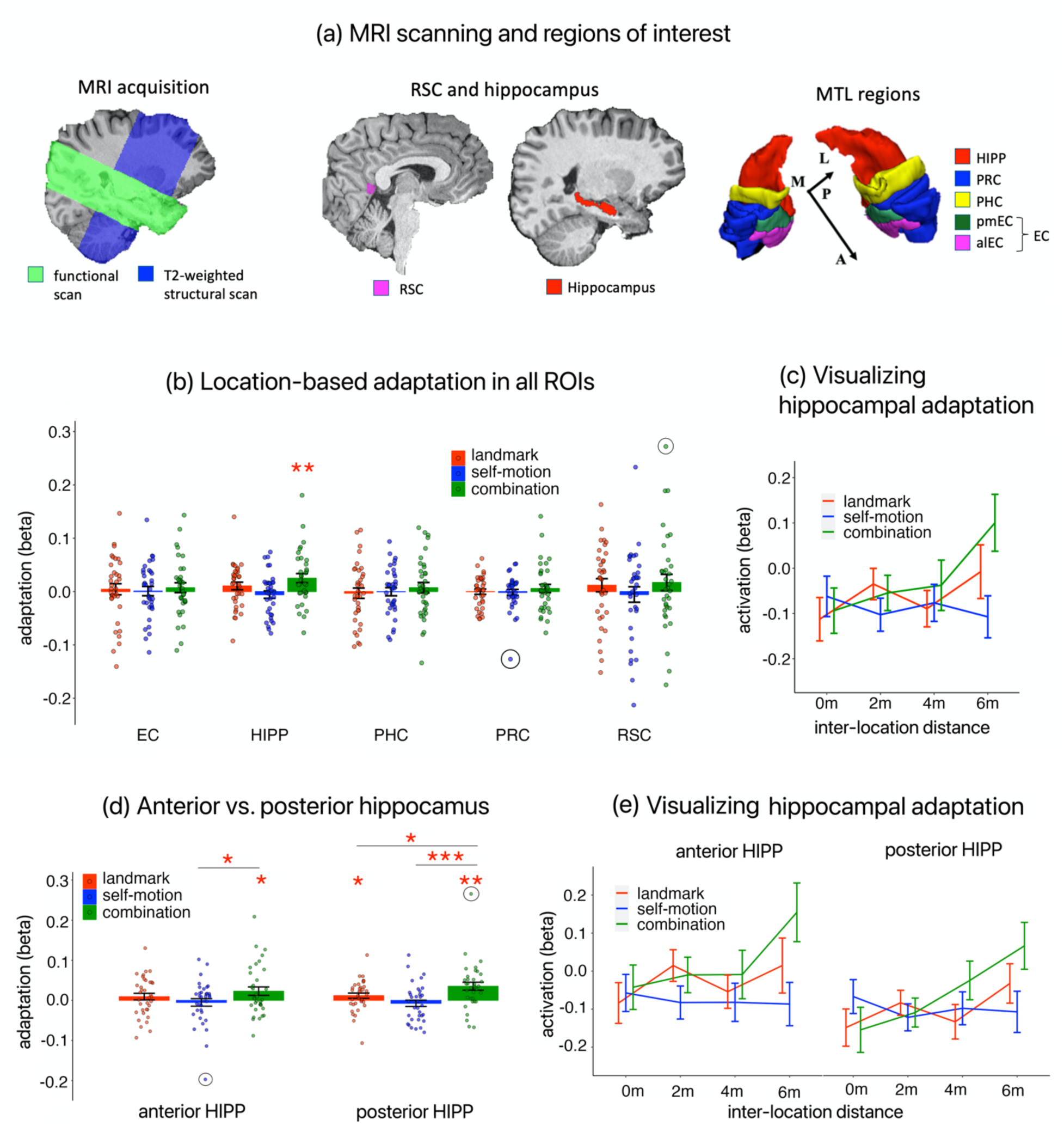
MRI acquisition, anatomical masks and location-based adaptation results. **(a)** MRI scans and regions of interest. Left: functional scan and T2-weighted structural scan overlaid on T1-weighted structual scan in one participant. Middle: anatomical masks of retrosplenial cortex (RSC) and hippocampus overlaid on T1-weighted structural scan in one participant,. Right: manually segmented anatomical masks of MTL regions in one participant. **(b)** Location-based adaptation across ROIs. Adaptation is indexed by beta weight for a parametric regressor modeling the modulatory effect of inter-location distance on BOLD responses. Dots represent individual participants. Dark circles indicate statistical outliers, which were winsorized to the nearest inlier for statistical tests. **(c)** Visualization of adaptation effects in the hippocampus. Hippocampal activation, which corresponds to beta weight estimated for the event regressor modeling the location occupation period, is plotted as a function of inter-location distance. **(d)** Location-based adaptation in the anterior and posterior hippocampus. **(e)** Visualization of adaptation effects in the anterior and posterior hippocampus. Dots represent individual participants. Dark circles highlight statistical outliers. Error bars represent ±SE. “*” – p_1-tailed_ < 0.05; “**” – p_1-tailed_ < 0.01; “***” – p_1-tailed_ < 0.001.

### Posterior Hippocampus Displays Stronger Adaptation in Combination Condition than both Single-Cue Conditions

The previous results showed that the hippocampus displayed location-based adaptation exclusively in the combination condition, reflecting a cue integration effect. However, although the hippocampal adaptation was numerically greater in the combination condition than both single-cue conditions, the difference was significant only when comparing the combination condition to the self-motion condition (t(34) = 2.686, p_1-tailed_ = 0.004, Cohen’s d = 0.481, BF_10_ = 10.947; a directional test was adopted because we expected stronger coding in the combination condition than single-cue conditions) but not to the landmark condition (t(34) = 1.285, p_1-tailed_ = 0.104, Cohen’s d = 0.217, BF_10_ = 0.686).

The lack of a statistically significant difference between the combination and landmark conditions raises an alternative explanation for the earlier findings. Specifically, the exclusive presence of hippocampal adaptation in the combination condition may not reflect genuine cue integration. Instead, it could be due to the use of an allocentric strategy based on the landmark in both the combination and landmark conditions, as opposed to an egocentric strategy based on self-motion cues in the self-motion condition.

To test this alternative explanation, we divided the hippocampus into anterior and posterior portions (Methods). Compared to the anterior part, the posterior hippocampus is more involved in spatial navigation (Grady, 2020). Furthermore, the precision of spatial representations is higher in the posterior hippocampus (Brunec et al., 2018; Evensmoen et al., 2015; Kjelstrup et al., 2008), aligning with the notion that cue integration enhances the precision of neural representations (Cheng et al., 2007). Therefore, we hypothesized that the cue integration effect would be more readily detectable in the posterior hippocampus.

Focusing on the posterior hippocampus (Figure 3d), we found that location-based adaptation was significantly greater than zero in the combination condition (t(34) = 3.412, p_1-tailed_ < 0.001, BF_10_ = 58.433; after winsorizing one outlier, t(34) = 4.065, p_1-tailed_ < 0.001, BF_10_ = 205.392). This adaptation was also significantly greater than that in the landmark condition (t(34) = 2.063, p_1-tailed_ = 0.023, BF_10_ = 2.311) and the self-motion condition (t(34) = 3.401, p_1-tailed_ < 0.001, BF_10_ = 39.053; after winsorizing one outlier, t(34) = 3.794, p_1-tailed_ < 0.001, BF_10_ = 102.828). Location-based adaptation in the landmark condition was also significantly greater than zero (t(34) = 1.787, p_1-tailed_ = 0.041, BF_10_ = 1.446). These results indicate a genuine cue integration effect in the posterior hippocampus.

In the anterior hippocampus (Figure 3d), location-based adaptation was significantly greater than zero in the combination condition (t(34) = 2.179, p_1-tailed_ = 0.018, BF_10_ = 2.855). This adaptation was significantly greater than that in the self-motion condition (t(34) = 2.850, p_1-tailed_ = 0.037, BF_10_ = 1.602; after winsorizing one outlier, t(34) = 1.899, p_1-tailed_ = 0.033, BF_10_ = 1.741). However, adaptation in the combination condition was not significantly greater than that in the landmark condition (t(34) = 1.061, p_1-tailed_ = 0.148, BF_10_ = 0.514).

In summary, compared to the anterior hippocampus, we found that the cue integration effect was more readily detectable in the posterior hippocampus, which displayed stronger positional coding in the combination condition than in both single-cue conditions. Intriguingly, we also detected relatively weak positional coding in the landmark condition in the posterior hippocampus. This finding suggests that the hippocampus contained spatial information for landmarks alone, potentially reflecting an inclination toward allocentric navigation. However this coding was significantly smaller than the coding in the combination condition. Thus, the enhanced positional coding in the combination condition compared to single-cue conditions can not be explained by participants adopting different navigation strategies associated with different cue types (i.e., an allocentric strategy with landmarks and an egocentric strategy with self-motion cues) but instead reflects true integration of spatial cues.

### Integrators, but not Non-integrators, Display Hippocampal Positional Coding Reflecting Cue Integration

The preceding results have highlighted hippocapmus’s unique role in representing spatial information derived from converging spatial inputs. However, it remains to be determined whether the hippocampus actively facilitated cue integration, leading to improved behavioral performance. We categorized participants based on whether they exhibited a cue integration effect behaviorally, defined as greater behavioral accuracy in the combination condition compared to both single-cue conditions.

*<u>Behavioral differences</u>*. Participants were divided into two groups: integrators (n = 24), who demonstrated higher mean accuracy averaged across test locations in the combination condition than in each single-cue condition, and non-integrators (n = 11), who did not.

As shown in Figure 4a (left panel), non-integrators performed better than integrators in the landmark condition (mean ACC = 0.786 vs. 0.699, t(33) = 2.318, p = 0.027, Cohen’s d = 0.974, BF_10_ = 2.438). Non-integrators’ performance in the combination condition was nearly identical to the landmark condition (mean ACC = 0.788 vs. 0.786, t(10) = 0.773, p = 0.457, Cohen’s d = 0.233, BF_10_ = 0.383). Notably, integrators and non-integrators exhibited similar behavioral performance in the combination condition (mean ACC = 0.773 vs. 0.788, t(33) = 0.465, p = 0.645, BF_10_ = 0.373).

Representational uncertainty, as estimated through the extended signal detection model (Methods – Cognitive modeling: Extended signal detection theory), showed results consistent with behavioral accuracy (Figure 4c). For integrators, representational uncertainty in the combination condition was lower than the best single-cue condition (t(23) = -5.213, p < 0.001, BF_10_ = 865). In contrast, non-integrators’ representational uncertainty did not differ between the combination and best single cue conditions (t(10) = 2.041, p = 0.069, BF_10_ = 1.347), and was numerically larger than in the landmark condition (mean = 0.867 m vs. 0.789 m). They exhibited lower representational uncertainty than integrators in the landmark condition (t(33) = -2.809, p = 0.008, BF_10_ = 5.776) and the best single-cue condition (t(33) = -2.549, p = 0.016, BF_10_ = 3.599). The two groups did not differ in representational uncertainty in the combination condition (t(33) = 0.444, p = 0.660, BF_10_ = 0.370).

To gain a deeper understanding of how integrators and non-integrators processed landmarks and self-motion cues in the combination condition, we conducted an additional modeling analysis (see Methods – Cognitive modeling: Comparing cue-handling strategies in combination condition). We evaluated four different cue-handling strategies:

1. **Bayesian cue integration.** Participants combined landmark and self-motion estimates in a statistically optimal way, weighting each by its reliability to maximize positional precision. This model is also referred to as the maximum-likelihood estimation (MLE) model (Cheng et al., 2007).
2. **Bayesian cue alternation.** Participants alternated between landmarks and self-motion cues across trials, with alternation proportions determined by relative cue reliabilities.
3. **Landmark dominance.** Participants relied exclusively on landmarks, ignoring self-motion cues.
4. **Self-motion dominance.** Participants relied solely on self-motion cues, ignoring landmarks.

Unlike the prior modeling analysis, where the precision and bias parameters for the combination condition were allowed to vary freely (Figure 2b, Figure 4c), here these parameters were derived from those in the single-cue conditions, according to the cue-handling strategy. For each participant and strategy, we computed the Akaike Information Criterion (AIC) and summed those values across participants, with lower AIC indicating a better fit. Models incorporating sensory biases were compared to those without such biases.

For integrators (Figure 4d), the Bayesian cue integration model assumming no sensory biases provided the best fit. The second best-fitting model, the landmark dominance model assuming no sensory biases, received little support compared to the Bayesian cue integration model (delta AIC = 47 > 10) (Burnham & Anderson, 2002). Consistently, integrators’ mean representational uncertainty, averaged across test locations, in the combination condition was significantly lower than the best single-cue condition (t(23) = -5.213, p < 0.001, BF_10_ = 865.257) and did not differ from the MLE prediction (t(23) = 1.023, p = 0.317, BF_10_ = 0.343), indicating statistically optimal cue integration (Figure 4c). In contrast, for non-integrators (Figure 4e), the landmark dominance model assuming no sensory biases was the winning model. The second best-fitting model was the cue alternation model assuming no sensory biases (delta AIC = 4). Consistently, non-integrators’ mean representational uncertainty in the combination condition was significantly larger than the MLE prediction (t(10) = 5.810, p < 0.001, BF_10_ = 182;) and did not significantly differ from the best single-cue condition (t(10) = 2.041, p = 0.069, BF_10_ = 1.347), which corresponded to the landmark condition (Figure 4c). These results indicate lack of cue integration among non-integrators. In sum, integrators and non-integrators adopted distinct strategies in the combination condition: while integrators predominantly used the Bayesian cue integration strategy, non-integrators primarily used the landmark dominance strategy.

**Figure 4.**
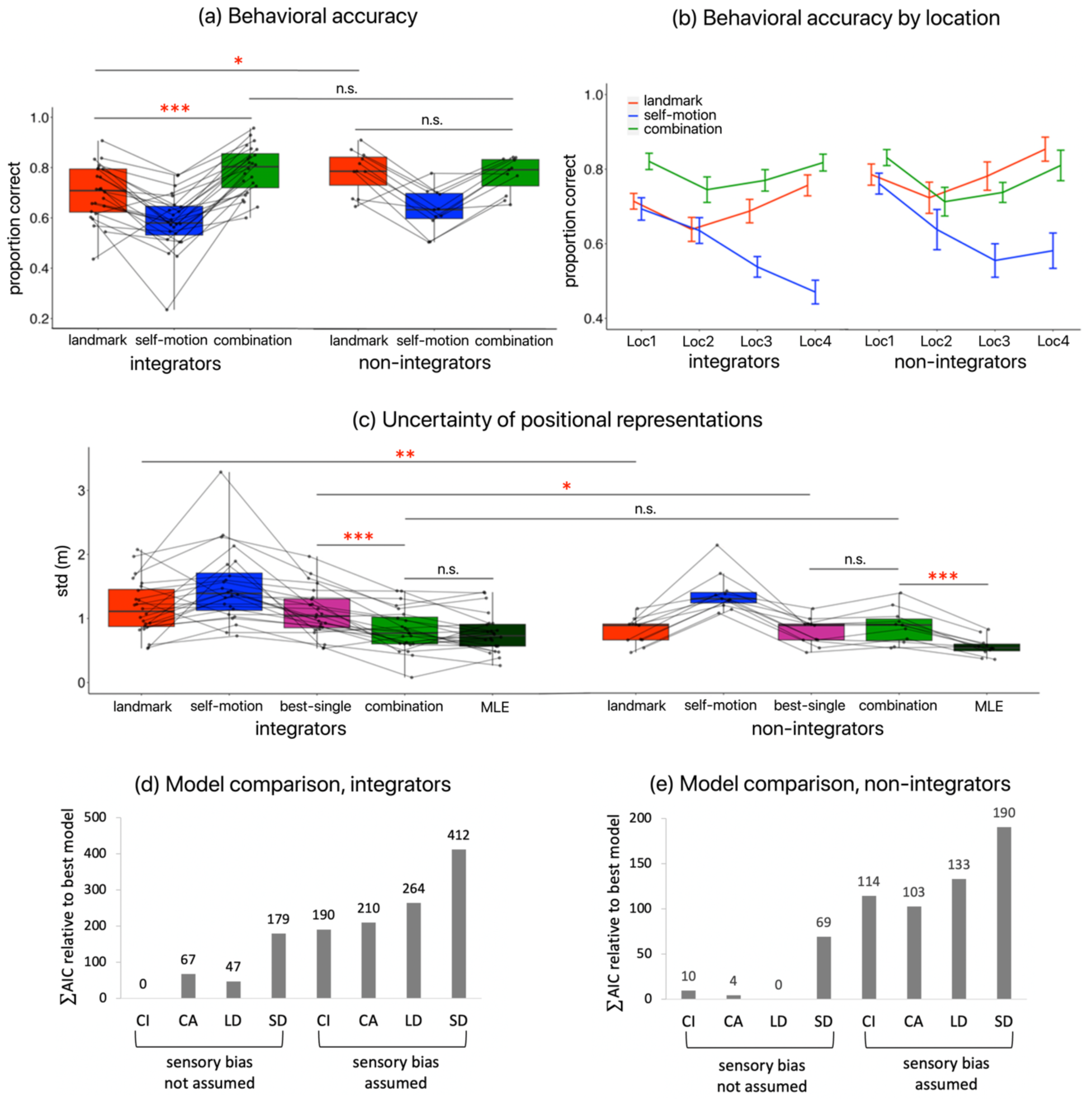
Behavioral differences between integrators and non-integrators. (a) Boxplots of behavioral accuracy by cue condition, shown separately for integrators and non-integrators. (b) Behavioral accuracy plotted by cue type and test location, for each group. (c) Boxplots of representational uncertainty from cognitive modeling, including the “best_single” uncertainty and MLE prediction. The MLE model corresponds to the Bayesian cue integration (CI) model in (d) and (e). (d) Results of model comparison for integrators. “CI” – Bayesian cue integration, “CA” – Bayesian cue alternation, “LD” – landmark dominance, “SD” – self-motion dominance. The Bayesian cue integration (CI) model corresponds to the MLE model in (c). (e) Results of model comparison for non-integrators, structured as in (d). Dots represent individual participants. Error bars represent ±SE. “*” – p < 0.05, “**” – p < 0.01, “***” – p < 0.01.

*<u>Differences in location-based adaptation.</u>* As shown in Figure 5a, integrators exhibited significant hippocampal adaptation in the combination condition (t(23) = 2.885, p_1-tailed_ = 0.004, Cohen’s d = 0.589), and the Bayes factor indicates strong evidence for the alternative hypothesis (BF10 = 11.202 > 10). The hippocampus exhibited no significant adaptation effects in the two single-cue conditions (landmark, t(23) = 0.980, p_1-tailed_ = 0.169, BF_10_ = 0.542; self-motion, t(23) = -0.817, p_1-tailed_ = 0.789, BF_10_ = 0.129), and Bayes factors indicate evidence favoring the null hypothesis (BF_s10_ < 1). No significant adaptation effects were detected in other ROIs (ts < 1.35, ps > 0.09, Cohen’s ds < 0.28, BF_s10_ < 0.86).

In contrast, non-integrators displayed no significant hippocampal adaptation in the combination condition (t(10) = 0.773, p_1-tailed_ = 0.229, Cohen’s d = 0.233; Figure 5b), with the Bayes factor indicating evidence favoring the null hypothesis (BF10 = 0.578 < 1). Considering the difference in sample size between the two groups, we conducted the power analysis. This analysis confirmed that the absence of hippocampal adaptation in the combination condition in non-integrators was not merely due to smaller sample size: 116 participants would be needed in the non-integration group to achieve a 80% power, versus only 20 in the integration group. Additionally, non-integrators exhibited no significant adaptation effects were observed in the single-cue conditions in the hippocampus either (landmark, t(10) = 1.096, p_1-tailed_ = 0.149, Cohen’s d = 0.330, BF_10_ = 0.811; self-motion, t(10) = -0.244, p_1-tailed_ = 0.596, Cohen’s d = -0.074, BF_10_ = 0.252), nor in other ROIs in any conditions (ts < 1.7, ps > 0.05, BF_s10_ < 1.6).

For integrators, hippocampal adaptation in the combination condition was significantly greater than in the self-motion condition (t(23) = 2.600, p_1-tailed_ = 0.005, Cohen’s d = 0.576, BF_10_ = 9.917), but was only marginally greater than in the landmark condition (t(23) = 1.629, p_1-tailed_ = 0.051, Cohen’s d = 0.348, BF_10_ = 1.416). Therefore, similar to the previous analyses of all participants, we conducted more detailed analyses by examining the anterior hippocampus and the posterior hippocampus separately. The results were similar to those for all participants (Figure 3d). Focusing on the posterior hippocampus (Figure 5c), we found significant location-based adaptation in the combination condition (t(23) = 3.057, p_1-tailed_ = 0.003, BF_10_ = 15.792; after winsorizing one outlier, t(23) = 3.551, p_1-tailed_ < 0.001, BF_10_ = 44.067). This adaptation was significantly greater than that in the landmark condition (t(23) = 2.179, p_1-tailed_ = 0.020, BF_10_ = 3.037) and the self-motion condition (t(23) = 3.133, p_1-tailed_ = 0.002, BF_10_ = 18.423; after winsorizing one outlier, t(23) = 3.609, p_1-tailed_ < 0.001, BF_10_ = 49.789). Regarding the anterior hippocampus, location-based adaptation was significantly greater than zero in the combination condition (t(23) = 2.241, p_1-tailed_ = 0.017, BF_10_ = 3.381), which was significantly stronger compared to the self-motion condition (t(23) = 1.939, p_1-tailed_ = 0.032, BF_10_ = 2.040; after winsorizing one outlier, t(23) = 2.049, p_1-tailed_ = 0.026, BF_10_ = 2.441) but not to the landmark condition (t(23) = 1.229, p_1-tailed_ = 0.116, BF_10_ = 0.733).

For non-integrators (Figure 5d), the only significant effect we observed was a significant location-based adaptation in the combination condition in the posterior hippocampus (t(10) = 1.925, p_1-tailed_ = 0.042, BF_10_ = 2.208). Crucially, this adaptation was not significantly greater compared to either the landmark condition (t(10) = 0.248, p_1-tailed_ = 0.405, BF_10_ = 0.360) or the self-motion condition (t(10) = 1.381, p_1-tailed_ = 0.099, BF_10_ = 1.122), indicating no evidence of of cue integration. Instead, the significant adaptation in the combination condition likely reflects spatial coding for landmarks alone, given that non-integrators primarily adopted the landmark dominance strategy in the combination condition (Figure 4e) and the posterior hippocampus exhibited positional coding in the landmark condition when all participants were analyzed (Figure 3d).

**Figure 5.**
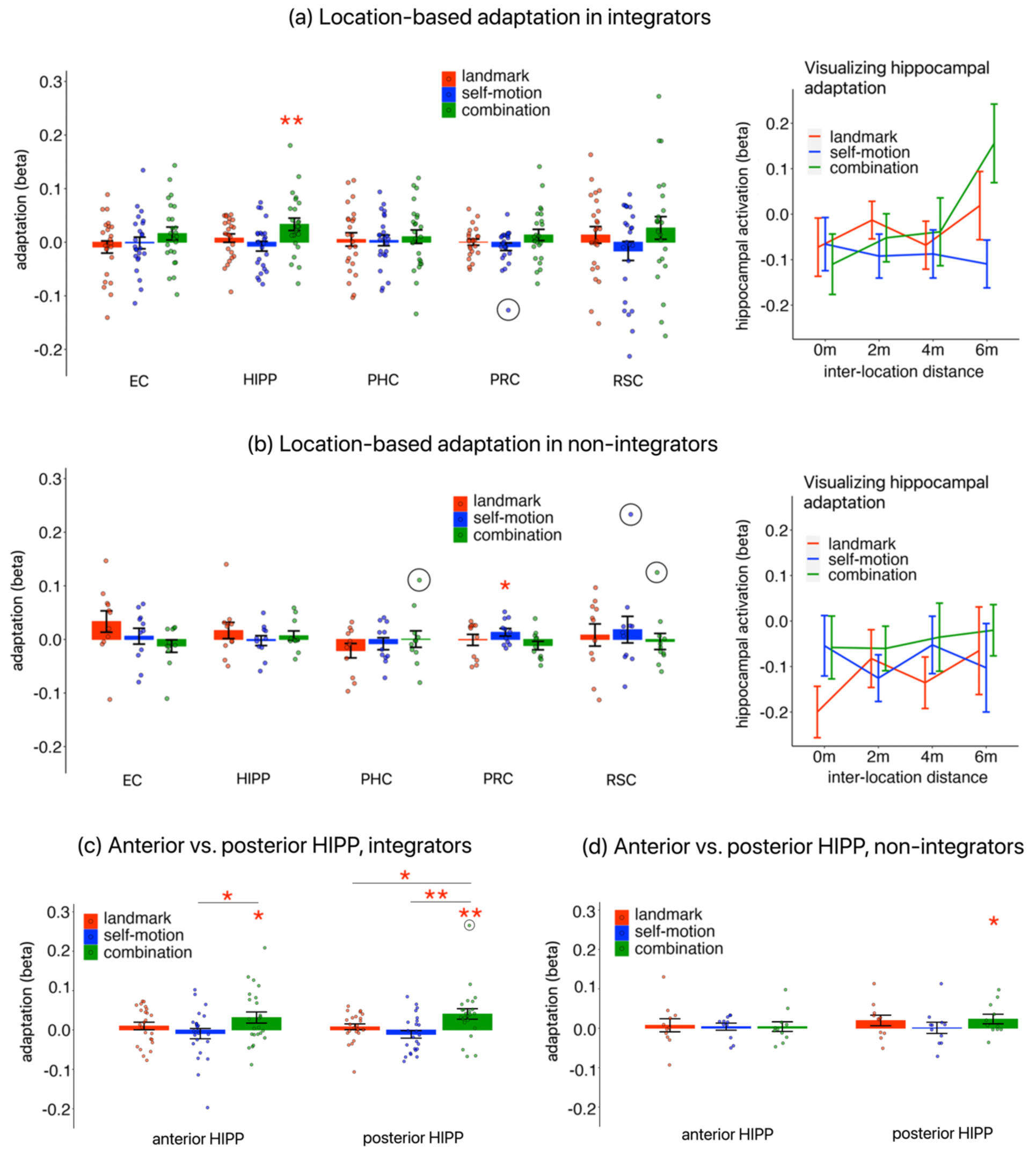
Differences between integrators and non-integrators in location-based adaptation. **(a)** Location-based adaptation in integrators. Left: adaptation by cue condition across ROIs. Right: hippocampal activation as a function of inter-location distance. **(b)** Location-based adaptation in non-integrators, structured as in (d). **(c)** Location-based adaptation in the anterior hippocampus and posterior hippocampus for integrators. **(d)** Location-based adaptation in the anterior hippocampus and posterior hippocampus for non-integrators. Dots represent individual participants. Dark circles indicate statistical outliers. Error bars represent ±SE. “*” – p_1-tailed_ < 0.05, “**” – p_1-tailed_ < 0.01.

*<u>Summary.</u>* The cue integration effect varied across participants, with roughly two-thirds showing behavioral evidence of integration—broadly consistent with prior work demonstrating inter-individual differences in cue integration (Zanchi et al., 2022; Zhao & Warren, 2015b). Non-integrators failed to integrate cues likely because they relied exclusively on landmarks in the combination condition, as landmarks alone supported strong performance.

Positional coding in the hippocampus was observed only in integrators and only in the combination condition, consistent with cue integration. The cue integration effect was particularly evident in the posterior hippocampus, which exhibited stronger positional coding in the combination condition compared to both single-cue conditions. In contrast, the cue integration effect was absent in the hippocampus for non-integrators, despite similar behavioral performance to integrators in the combination condition. These findings suggest that the hippocampal adaptation reflects benefit from cue integration, rather than overall task performance, highlighting an active role for the hippocampus—particularly its posterior portion—in integrating spatial cues. Although cue integration was more evident in the posterior hippocampus, the anterior hippocampus also exhibited neural activity indicative of cue integration—this region showed positional coding in the combination condition only, but not in any single-cue conditions. Therefore, in the following analyses, we analyzed the hippocampus as a whole without dividing it into anterior and posterior portions.

### Hippocampal Positional Coding Reflects Stimuli Rather than Behavioral Outputs

While the previous analyses established that the hippocampus was involved in spatial cue integration, its specific role in this process remains unclear. A key question is at what processing stage the hippocampus becomes engaged in cue integration. To address this question, we examined whether the hippocampal coding was more closely associated with the stimulus (i.e., true location) or the response (i.e., the participant’s self-reported location). If the hippocampus contributes at a relatively early stage of cue integration, its adaptation-based positional coding should be more related to true locations than self-reported locations. Conversely, stronger association with self-reported locations than true locations would suggest that the hippocampus gets involved at a relatively late stage. To assess whether the hippocampal coding reflected the stimulus or the response, we conducted neural space reconstruction and response-based adaptation analyses.

*<u>Neural space reconstruction</u>*. This analysis reconstructed the neural space with positional estimates for test locations, using neural distances derived from fMRI adaptation. Neural distance was quantified as activation to a location when preceded by another: lower activation indicates shorter neural distance between two successively visited locations. The neural space was then compared to the physical space. Data were aggregated across participants to maximize statistical power (Methods).

In the combination condition, the neural space reconstructed from hippocampal adaptation significantly resembled the physical space for all participants (p_1-tailed_ = 0.029; Figure 6a, left panels). Analyzing integrators revealed a similar result (p_1-tailed_ = 0.032; Figure 6a, middle panels). For non-integrators, the neural space did not significantly resemble the physical space (p_1-tailed_ = 0.145; Figure 6a, right panels), consistent with the lack of significant hippocampal adaptation for this group (Figure 5b). These findings suggest that hippocampal spatial coding reflected stimulus rather than behavior.

*<u>Response-based adaptation</u>.* To compute response-based adaptation, we constructed a general linear model in which the parametric modulation regressors were defined by participants’ reported locations rather than true locations. For instance, if a participant correctly identified Loc1 as Loc1 but misidentified Loc2 as Loc3, the parametric regressor had a value of |Loc1 – Loc3| = 4 m instead of |Loc1 – Loc2| = 2 m.

For all participants, the hippocampus did not show significant response-based adaptation in the combination condition, even at the uncorrected significance level (t(34) = 1.417, p_1-tailed_ = 0.083; Figure 6b, left panel); moreover, the Bayes factor indicates evidence favoring the null hypothesis (BF10 = 0.823 < 1). For integrators, the effect was marginally significant (t(23) = 1.715, p_1-tailed_ = 0.050, BF_10_ = 1.441). These results contrast the strong location-based hippocampal adaptation under the same condition (Figure 6b, right panel).

We wondered whether differences between location-based and response-based adaptation were due to differences in detection power, as the event sequences were designed based on true locations to maximize the detectability of adaptation effects (Methods). As expected, detection power was higher for true locations than for reported locations (t(34) = 6.163, p < 0.001, BF_10_ = 31365), but the magnitude of difference was negligible (mean = 48.5% vs. 47.1%), suggesting that detection power was unlikely to have confound the adaptation results.

**Figure 6.**
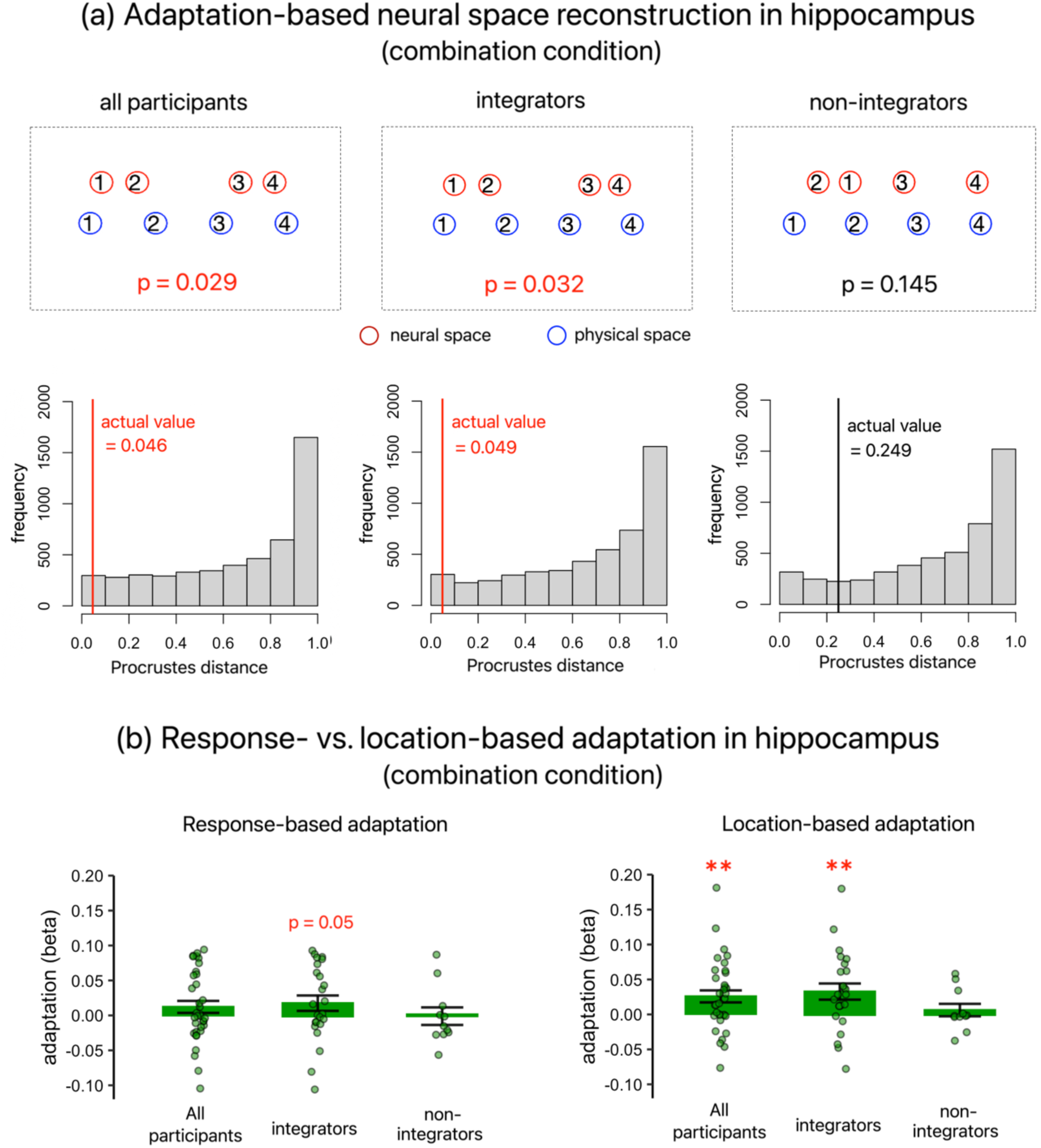
Comparing stimulus and response in hippocampal adaptation. **(a)** Adaptation-based neural space reconstruction in the hippocampus in the combination condition. Panels in the upper row depict neural space and physical space for all participants (left), integrators (middle), and non-integrators (right). Panels in the lower row show the actual Procrutes distance (vertical line) against the surrogate distribution (the histogram) for all participants (left), integrators (middle), and non-integrators (right). **(b)** Response-based adaptation in the hippocampus in the combination condition (left), compared to location-based adaptation (right, from Figure 3b, 5a, and 5b). Dots represent data from individual participants, and error bars represent ±SE. “**” – p_1-tailed_ < 0.01.

*<u>Summary</u>.* The neural space reconstruction and response-based adaptation analyses yielded convergent results: hippocampal adaptation in the combination condition was more closely tied to the stimulus than behavior. These findings suggest that the hippocampus becomes engaged at a relatively early stage of cue integration (e.g., sensory-perceptual processing), which already enhances the precision of positional representations.

### Hippocampus Serves as Functional Hub within MTL-RSC Network

The preceding results have established the hippocampus’ central role in spatial cue integration within the MTL-RSC network. This finding begs the question of why cue integration predominantly occurred in the hippocampus. We speculate the hippocampus might function as a network hub (van den Heuvel & Sporns, 2013), integrating information from multiple sources due to its dense interconnections with other MTL regions (Witter et al., 2000) and RSC (Vann et al., 2009).

We performed a graph analysis—an established methodology to characterize network connectivity features (Bullmore & Bassett, 2011)—using beta time-series extracted from the location occupation period. We calculated “betweenness centrality”— the number of shortest paths passing through the node—as an index of nodal hubness (Kong et al., 2017). The two hemispheres were analyed separately to maximize the number of nodes (Methods).

When group-level functional connectivity matrices were analyzed (Figure 7a), both left and right hippocampus were identified as hubs in all cue conditions (Figure 7b). No other regions were identified as hubs, with betweenness centrality equal to 0. The results remained consistent when we analyzed participant-specific functional connectivity matrices by submitting betweenness centrality to a repeated-measures ANOVA, with ROI, hemisphere, and cue condition as independent variables. The main effect of ROI was significant (F(5, 170) = 46.397, p < 0.001, 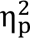 = 0.577): the hippocampus showed significantly greater betweenness centrality than all other ROIs (ts > 4.8, ps < 0.001). No other effects were significant (Fs < 2.1, ps > 0.12, 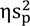 < 0.006).

**Figure 7.**
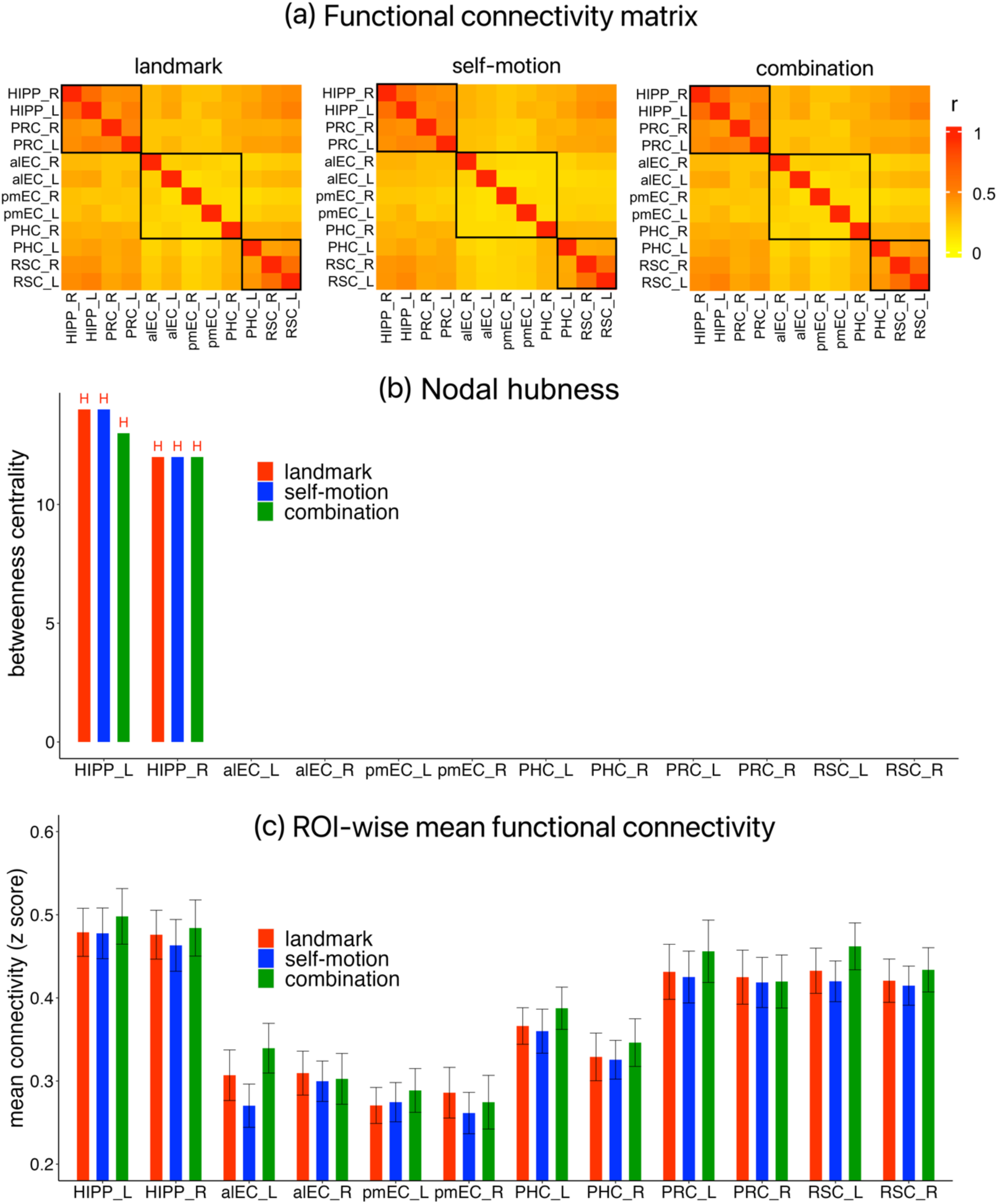
Results of task-based connectivity analyses. **(a)** Graph analysis, based on group-level functional connectivity matrices calculated from beta time-series. The modular structure of the MTL-RSC network is shown for the landmark (left), self-motion (middle), and combination condition (right). **(b)** Betweenness centrality, which indexes nodal hubness, displayed for each ROI and each cue condition. “H” indicates a hub. Results were derived from the group-level functional connectivity matrix thresholded at 60%. **(c)** Mean functional connectivity for all ROIs. Error bars represent ±SE.

To gain a more straightforward understanding of the functional connectivity pattern, we analyzed the mean connectivity strength (Figure 7c), calculated as the average of one region’s connectivity with all other regions. This measurement was submitted to a repeated-measures ANOVA, with ROI, hemisphere, and cue condition as independent variables. The main effect of ROI was significant (F(5, 170) = 107.161, p < 0.001, 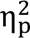 = 0.759): the hippocampus exhibited stronger mean connectivity than all other ROIs across cue conditions (ts > 4.4, ps < 0.001).

In summary, we found that the hippocampus served as a central node in the MTL-RSC network. This result aligns with our earlier finding that the hippocampus was the only region in this network to exhibit positional coding associated with cue integration.

### Cue-Integration-Related Positional Coding beyond ROIs

To explore positional coding beyond our predefined ROIs, we conducted the voxel-wise analysis of location-based adaptation, using a nonparametric permutation-based method for multiple comparisons correction across the entire search volume (Nichols & Holmes, 2002). We adopted a cluster-defining threshold of T > 3 for cluster-level inference. Full results are provided in Table S1.

When analyzing all participants (Figure 8a), we detected significant clusters with peaks in the angular gyrus, precuneus, middle temporal gyrus in the combination condition. No significant clusters (_psFWE-corr,1-tailed_ > 0.08) or voxels (_psFWE-corr, 1-tailed_ > 0.4) were found in either single-cue condition.

A similar pattern was observed for integrators (Figure 8b). We detected significant clusters with peaks in the dorsal posterior cingulate cortex, precuneus, middle temporal gyrus, and angular gyrus. No significant clusters (_psFWE-corr, 1-tailed_ > 0.3) or voxels (_psFWE-corr, 1-tailed_ > 0.7) were found in the two single-cue conditions.

In contrast, non-integrators showed no significant clusters (_psFWE-corr,1-tailed_ > 0.2) or voxels (_psFWE-corr,1-tailed_ > 0.7; Figure 8c) in the combination condition. However, in single-cue conditions, we detected one cluster whose peak resided in the superior occipital gyrus in the landmark condition, and three clusters with peaks in the precuneus, angular gyrus, and middle occipital gyrus in the self-motion condition.

In sum, we identified positional coding related to cue integration in the precuneus (likely POS), dorsal posterior cingulate cortex, and angular gyrus. Notably, integrators exhibited positional coding in the combination condition but not in single-cue conditions, whereas non-integrators exhibited the opposite pattern. These findings highlight inter-individual differences and implicate a broader network supporting spatial cue integration beyond the hippocampus.

**Figure 8.**
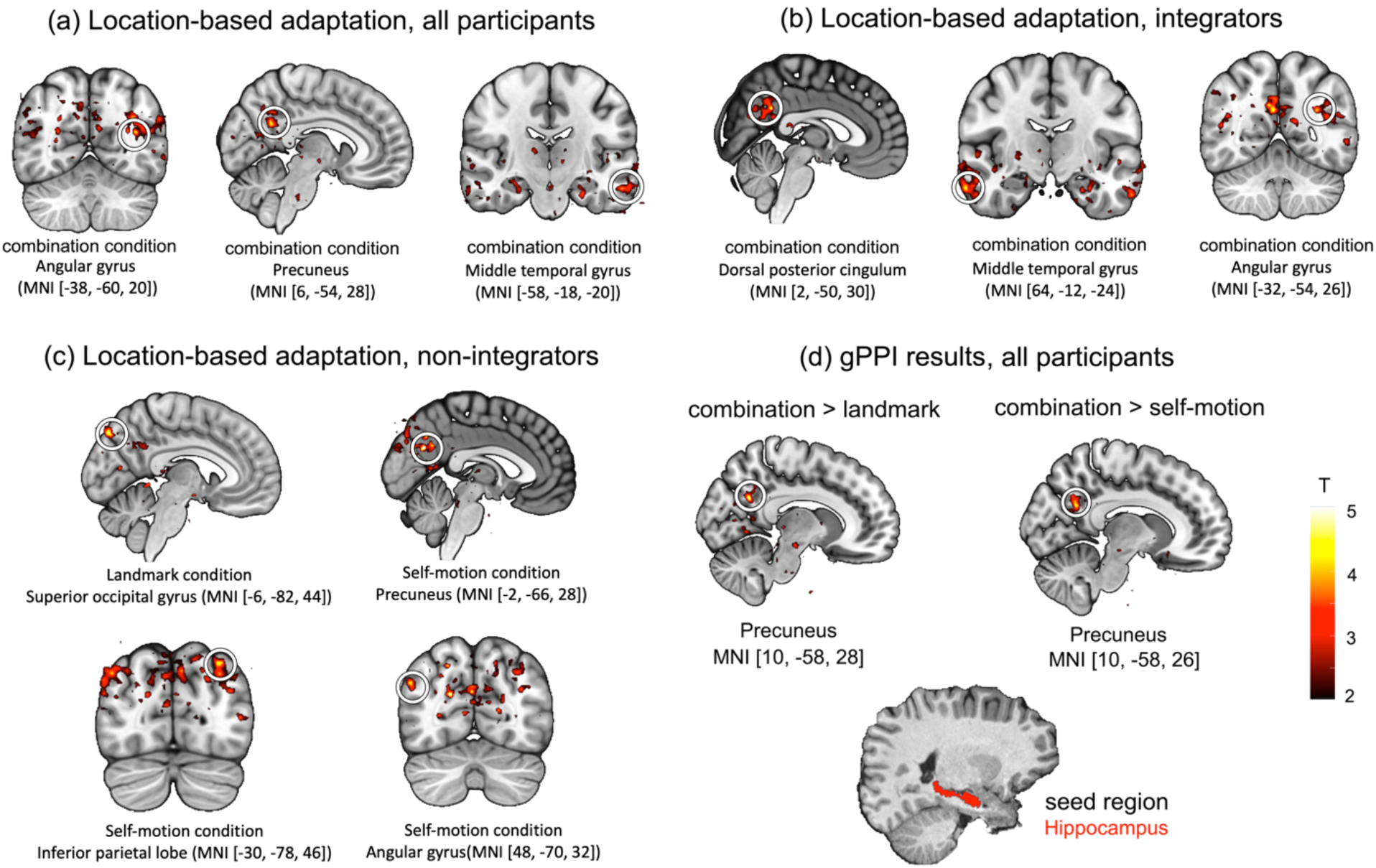
Spatial cue integration involves a broader network beyond hippocampus. **(a)** Voxel-wise analysis of location-based adaptation for all participants. **(b)** Same analysis for integrators only. **(c)** Same analysis for non-integrators only. **(d)** gPPI analysis for all participants using the bilateral hippocampus as the seed region. For illustration, results are displayed on the MNI template, thresholded at T > 2.

### Hippocampus-Precuneus Connectivity Contributes to Spatial Cue Integration

To explore the functional connectivity that contributed to the spatial cue integration effect observed in the hippocampus, we conducted the generalized psychophysiological interaction (gPPI) analysis, (McLaren et al., 2012), using the hippocampus as the seed region (Methods). Given the typically low statistical power of PPI analyses (O’Reilly et al., 2012), we performed this analysis only across all participants.

As shown in Figure 8d, we observed stronger hippocampal connectivity with the precuneus (likely POS) in the combination condition compared to the landmark condition (T = 5.82, _pFWE-corr,1-tailed_ = 0.007, MNI [10, -58, 28], voxel-level inference) and the self-motion condition (peak T = 4.87, MNI [10, -58, 26], k = 34, _pFWE-corr,1-tailed_ = 0.017, cluster-level inference). These results indicate that the hippocampus-precuneus interaction contributed to spatial cue integration. Notably, positional coding associated with cue integration was also detected in the precuneus (likely POS, peak voxel MNI [6, -54, 28], Figure 8a).

## Discussion

This study examined how the brain integrates diverse spatial cues to enhance navigation performance. Participants learned and retrieved spatial locations on a linear track while undergoing 3T fMRI scanning. Focusing on the MTL and RSC, we detected positional coding reflective of cue integration exclusively in the hippocampus. The cue integration effect was particularly pronounced in the posterior hippocampus. Spatial cue integration also recruited other areas such as the precuneus (putative POS). Our findings provide evidence for the unique hippocampal involvement in spatial cue integration and illuminate the intricate neural mechanisms underpinning this process.

We found that participants integrated landmarks and self-motion cues to improve navigation performance. Note that in all cue conditions, both cues were physically present, but single-cue conditions required participants to rely on a designated cue. This design minimized sensory differences across conditions, avoiding confounds in the fMRI results. However, in single-cue trials, participants might have struggled to ignore the task-irrelevant cue, disrupting their performance. This cue interference could create a spurious effect of cue integration, meaning that performance enhancement observed in the combination condition was due to the absence of cue interference rather than the benefit from cue integration.

We consider this explanation unlikely. Participants received intensive training prior to the MRI scanning. During scanning, they received explicit feedback defined by the task-relevant cue, reinforcing appropriate cue usage. Their adherence to instructions is supported by behavioral and modeling results (Figure 2a and Figure 2b, right panels): behavioral accuracy and representational precision increased as the test location neared the respective anchoring points—namely, the landmark in the landmark condition and the starting position of movement in the self-motion condition. These findings indicate that participants used the intended cues appropriately and that the enhanced performance in the combination condition reflects a genuine cue integration benefit rather than a lack of interference. This behavioral evidence of cue integration lays the foundation for investigating the underlying neural mechanisms using fMRI data.

Within the MTL-RSC network, the hippocampus was the only region to exhibit positional coding indicative of cue integration. This effect manifested as location-based adaptation that occurred exclusively in the combination condition. Notably, the posterior hippocampus showed particularly pronounced cue integration effect, displaying greater adaptation in the combination condition compared to both single-cue conditions. This finding aligns with its established role in in spatial navigation (Grady, 2020) and its capacity for high-precision spatial representations (Brunec et al., 2018; Evensmoen et al., 2015; Kjelstrup et al., 2008). Together, these findings suggest that combining cues sharpens neural representations of locations in the hippocampus, consistent with a core principle of cue integration (Cheng et al., 2007). Supporting this interpretation, the modeling results revealed enhanced representational precision in the combination condition compared to each single-cue condition.

Crucially, hippocampal positional coding emerged only in integrators who behaviorally benefited from cue integration, but not non-integrators, despite similar behavioral performance in the combination condition. These findings indicate that the hippocampal coding reflects performance enhancement brought by cue integration rather than absolute task performance. This finding also counters the abovementioned cue interference account: although cue interference was absent in the combination condition for both integrators and non-integrators, the hippocampal coding was absent for non-integrators. Thus, the mere absence of cue interference was insufficient to elicit integration-related hippocampal coding.

The lack of a cue integration effect in non-integrators may be partly explained by the principle of inverse effectiveness (Stein & Stanford, 2008), which posits that when a single cue already yields strong performance, adding another cue offers limited additional benefit. Although this principle still implies cue integration, it predicts limited benefit in uncertainty reduction. Such patterns have been observed in behavioral studies of cue integration (Chen et al., 2017, Exp. 1A, rich environment condition; Zanchi et al., 2022). However, this explanation alone is insufficient, as the MLE prediction of response variability was significantly lower than the observed variability in the combination condition, suggesting that uncertainty reduction was still possible. Cognitive modeling demonstrated that non-integrators relied excusively on landmarks and disregarded self-motion cues in the combination condition. This finding suggests that the limited perceived benefit of cue integration may have led non-integrators to adopt a heuristic strategy (i.e., the landmark dominance strategy) (Figure 4e). In this strategy, landmarks received full weight (= 1), exceeding the value predicted by cue relative reliabilities (< 1). For these participants, to elicit cue integration effect in behavior and thus integration-related positional coding in the hippocampus, a more challenging landmark condition may be necessary (see Chen et al., 2017, Exp. 1A, poor environment condition).

Our previous fMRI studies investigating single-cue navigation also support the idea that the hippocampus plays a unique role in spatial cue integration (Chen et al., 2019, 2024). These studies employed similar tasks and analyses and focused on the same ROIs. Participants relied on either landmarks or self-motion cues alone for spatial localization. Significant adaptation-based positional coding was obtained in EC (Chen et al., 2019) and RSC (Chen et al., 2024), but not in the hippocampus. To complement the adaptation approach, we later employed the representational similarity analysis (Epstein & Morgan, 2012), which revealed positional coding in the hippocampus as well as in RSC and PHC (Chen et al., 2025). Collectively, these findings suggest that spatial representation is not a unique or primary function of the hippocampus during navigation. Instead, the hippocampus appears to contribute specifically to spatial cue integration. The hippocampus’s unique involvement in spatial cue integration may inform the long-standing theoretical debates regarding its functional role, particularly in relation to the cognitive map, episodic memory, and relational memory theories. The cognitive map theory posits that the hippocampus constructs allocentric spatial representations that support flexible navigation, such as short-cutting (O’Keefe & Nadel, 1978). Our findings offer partial support for this theory: while the hippocampus does code for navigational space, its core function may lie in integrating diverse spatial information to support precise, memory-based navigation. This function extends beyond simple spatial representation and distinguishes the hippocampus from other MTL regions and RSC.

The episodic memory theory posits that the hippocampus binds diverse elements in an event into coherent memory traces (Scoville & Milner, 1957; Squire et al., 2004; Yonelinas et al., 2019). Spatial (where) and temporal (when) contexts are particularly central elements. We propose that this binding process can be conceptualized through the principle of probability multiplication, whereby combining multiple uncertain elements multiplicatively yields more precise event representations (Ekstrom & Yonelinas, 2020). This principle is used to model cue integration in behavioral research (Bromiley, 2013; Chen et al., 2025; McNamara & Chen, 2022) (see Figure 9 for concrete examples). In this light, our finding of hippocampal involvement in spatial cue integration computationally aligns with its established role in episodic memory.

**Figure 9.**
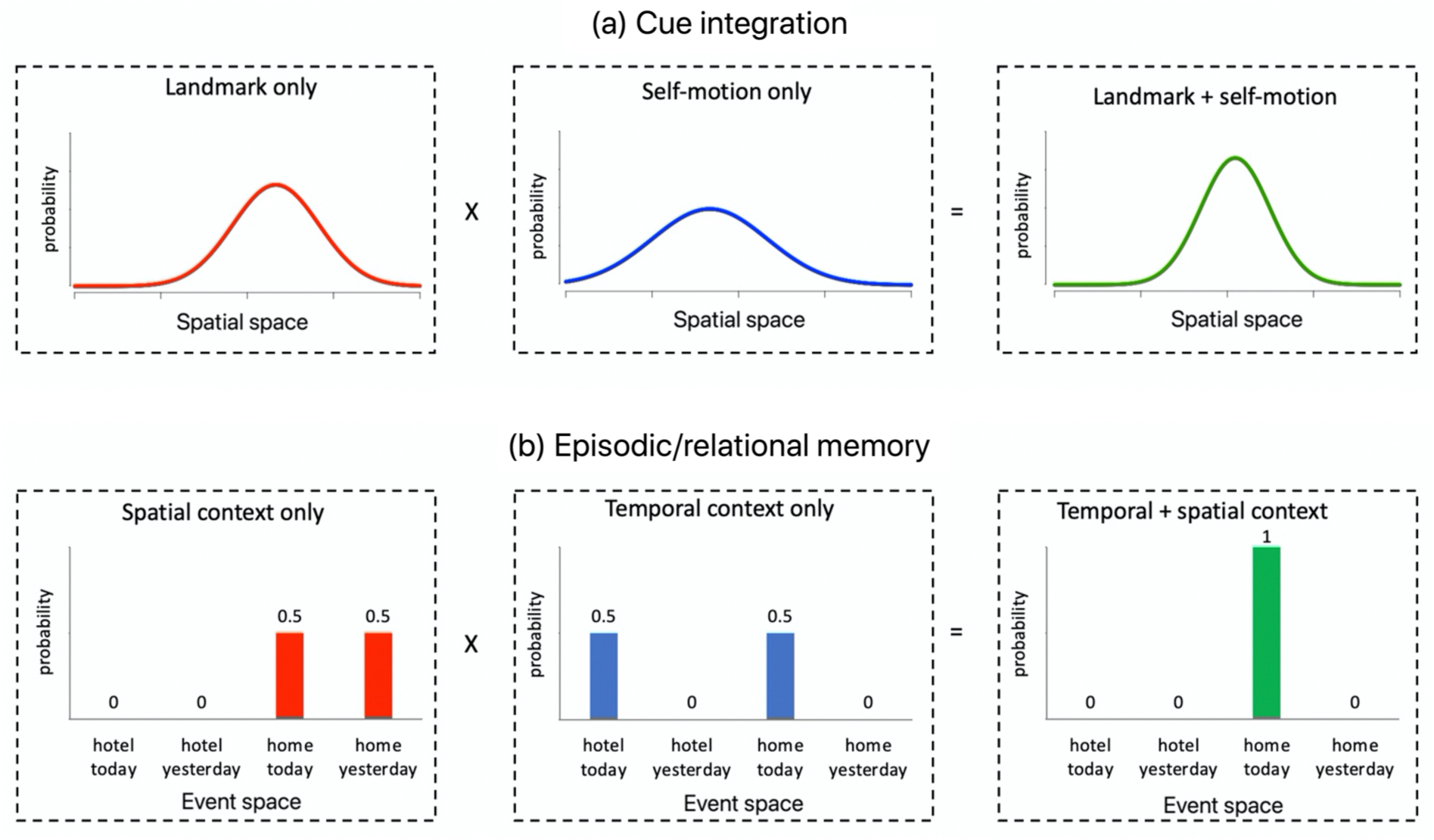
Computational parallel between cue integration and episodic/relational memory. Cue integration and episodic/relational memory may rely on the same principle of probabilistic multiplication to increase representational precision. **(a)** In cue integration, likelihood distributions from landmark-only (red) and self-motion-only (blue) cues are combined multiplicatively to produce a sharper joint distribution (green). Each likelihood distribution indicates the probability of each possible position being the true target location given the current spatial input. Single-cue likelihood distributions are assumed to be normal distributions. According to the basic probability tenet that the joint probability of two independent events occurring simultaneously is equal to the product of their individual probabilities, the joint likelihood distribution (the green curve) is obtained by multiplying the two single-cue likelihood distributions point-by-point. After normalization, the joint likelihood distribution is also a normal distribution but has higher precision than both single-cue distributions. **(b)** In episodic memory, combining multiple event elements yields more precise identification of the correct event. In this example, temporal context or spatial context alone yields uncertain event estimates, but combining their likelihood distributions multiplicatively also results in a more precise joint distribution that better disambiguates an event (“I ate breakfast at home today”) from similar alternatives. This framework can be extended to incorporate other discrete stimulus features involved in episodic and relational memory, such as color, location, person, object, etc.

The relational memory theory emphasizes the hippocampus’s role in encoding and retrieving associations among elements – for example, linking names with faces (Cohen & Eichenbaum, 1993). Episodic memory can be viewed as a specialized form of relational memory, enriched with spatiotemporal context. Therefore, the act of associating elements can also be conceptualized through the principle of probability multiplication (Figure 9), as this principle is invariant to the nature of the elements and thus applicable across a variety of stimulus types—for example, discrete vs. continuous, or spatialotemporal context vs. objects/persons. From this perspective, the hippocampus’s core function may lie in integrating diverse information sources to construct memory representations that are more precise than those based on individual elements alone. This integrative mechanism may constitute a common computational foundation across seemingly distinct cognitive domains, including cue integration, episodic memory, and relational memory.

In addition to hippocampal involvement in spatial cue integration, the present study also offers deeper insights into how spatial cue integration may be implemented in the hippocampus. We found that hippocampal adaptation-based positional coding was stimulus-driven rather than behavior-driven. This finding aligns with previous human fMRI studies showing that fMRI adaptation is sensitive to physical properties of stimuli (Chen et al., 2024; Epstein & Morgan, 2012). Such stimulus-driven coding suggests that the hippocampus participates at a relatively early stage of the cue integration process. We speculate that the hippocampus integrates sensory-perceptual inputs from multiple spatial cues, for example, processed spatial estimates based on individual cues. The hippocampus may contain a class of location-tuned neural units, which are oriented towards sensory-perceptual processing and exhibit adaptation properties. These units may achieve spatial cue integration either at the single-unit or population level. At the single-unit level, input from a single spatial cue may be insufficient to elicit robust activity in these neural units. Activating these neural units likely requires convergent inputs from multiple cues, possibly via a coincidence detection mechanism, as has been observed in hippocampal place cells (Jarsky et al., 2005; Takahashi & Magee, 2009). At the population level, theoretical models suggest that cue integration can be realized through probabilistic population codes, a way the brain represents uncertainty about sensory information using the collective activity of a population of neurons. Individual cues are represented as separate probabilistic population codes, which can simply be summed to produce a joint probabilistic code that is more precise than any individual cue (Ma et al., 2006).

The hippocampus’ role in spatial cue integration might be supported by its connectivity pattern. Within the MTL-RSC network, the hippocampus was identified as the sole hub. This finding aligns with a previous study based on resting-state fMRI data (Shah et al., 2018). This consistency between task-based and resting-state connectivity suggests that the hippocampus inherently occupies an privileged position for integrating various sources of neural information (van den Heuvel & Sporns, 2013). Beyond the MTL-RSC network, the hippocampus exhibited stronger connectivity with precuneus (likely POS) in the combination condition compared to each single-cue condition. Information exchange between the hippocampus and POS is crucial for forming spatial memories (Diersch et al., 2021). Notably, POS also exhibited cue-integration-related positional coding. These findings suggest that, at a larger scale, spatial cue integration recruits a decentralized system. This system involves multiple regions working collaboratively to combine various spatial inputs, thereby enhancing the flexibility and robustness of cue integration (Zhang et al., 2016).

Non-human studies have extensively explored how spatial cues interact during navigation, but fail to provide conclusive evidence about the neural basis of spatial cue integration, due to the limitations outlined in the introduction. While our study overcomes these limitations, it has constraints. First, we used visual optic flow instead of body-based self-motion cues to stimulate path integration, as participants have to remain motionless during MRI scanning. Consequently, caution is advised when comparing our study to non-human animal studies. Second, we obtained little evidence of positional coding in single-cue conditions. Single-cue conditions were designed to be challenging, which is necessary for the emergence of cue integration effect (Stein & Stanford, 2008). Additionally, we used 3T fMRI, which produced BOLD signals with lower intensity compared to 7T fMRI in our previous investigations of single-cue navigation (Chen et al., 2019, 2024). It is also possible that positional coding for individual cues might manifest in a different format (Figure S6). Finally, our study could not determine whether spatial cue integration occurred in the egocentric or allocentric reference frame (Negen et al., 2021). The starting position of movement was not randomized, preventing disentanglement of the test location’s allocentric position from the length of the path leading to it (Chen et al., 2024).

## Conclusions

The current study examines a fundamental question in spatial navigation—how the brain integrates landmarks and self-motion cues to enhance navigation performance. Our findings reveal that the hippocampus plays a distinct role in this integration process, offering critical insights into its underlying neural mechanisms. More broadly, these findings help reconcile the long-standing debate over whether the hippocampus primarily supports spatial cognition and episodic/relational memory, suggesting that its core function is to integrate diverse sources of information to enable precise long-term memory.

## Abbreviations

RSC: Retrosplenial cortex
MTL: Medial temporal lobe
HIPP: Hippocampus
PHC: Parahippocampal cortex
PRC: Perirhinal cortex
EC: Entorhinal cortex
alEC: Anterior-lateral entorhinal cortex
pmEC: Posterior-medial entorhinal cortex
POS: Parieto-occipital sulcus

## Acknowledgements

This research was supported by the National Natural Science Foundation of China (NSFC; #32100839), STI 2030 - Major Projects (#2021ZD0200409), and Cao Guangbiao High Science and Technology Foundation, Zhejiang University (ZJU; 2020QN002). We thank Timothy P. McNamara and Russel A. Epstein for their valuable feedback on the manuscript. We are also grateful to Qiuping Ding from the Center for Brain Imaging Science and Technology, Zhejiang University, for technical assistance with the MRI scanning, and to Yingyan Chen (陈颖妍) and Xiao Yu (禹潇) for their help with data collection.

## Declaration of generative AI and AI-assisted technologies in the writing process

During the preparation of this work the author(s) used ChatGPT in order to improve the language and readability of the manuscript, with caution. After using this tool, the authors reviewed and edited the content as needed and take full responsibility for the content of the publication.

## Methods

### Participants

We recruited thirty-eight healthy young adults from Zhejiang University (20 males, age range: 19 to 30 years; mean age = 23.8). All participants were right-handed, had normal or corrected-to-normal vision, and reported no history of neurological diseases. Three participants were excluded from the analysis due to dropout or technical issues that compromised the fMRI data quality. Written informed consent was obtained from all participants prior to the experiment, and they received monetary compensation for their participation. The study was approved by the local ethics committee of Zhejiang University, Hangzhou, China.

### Stimuli and navigation task

The virtual reality (VR) environment for the tasks was developed using Worldviz 5.0 software (https://www.worldviz.com). The environment featured a central linear track marked by a red arrow and a tree. The red arrow was positioned at the imagery midline of the linear track, and the tree was positioned slightly off the midline by 0.5 vm. Four differently colored balls were positioned at four test locations in the imagery midline of the linear track, between the red arrow and the tree. The four test locations were equally spaced, with 2 meters intervals between adjacent locations (Figure 1a).

*<u>Learning task</u>*. Prior to the MRI scanning, participants underwent a training session on a separate day (see the video “task_demo.mp4” – learning). They were required to memorize the locations of the four balls situated at the four equally-spaced test locations (Figure 1a). Initially, they were instructed to memorize the colors of the balls. Subsequently, they navigated along the linear track, utilizing both self-motion cues and the visible landmark (i.e., the tree), to learn the specific positions of each ball separately. Each ball was learned twice, with the presentation order counterbalanced across the four balls.

*<u>Test task (fMRI task): Location identification task.</u>* Participants performed the location identification task while undergoing functional MRI scanning (see the video “task_demo.mp4” – test). As shown in Figure 1b, during the location identification task, participants were passively transported to one of the four test locations, where they were asked to recall the color of the ball positioned at the test location. All the balls remained invisible throughout. The time course of the trial is as follows. First, the participant was passively transported from the red arrow to the test location. The tree was not displayed during the movement, and briefly displayed for 0.7 seconds after participant had arrived at the test location. The purpose of this manipulation was to minimize the overlap of low-level sensory inputs between different cue conditions as much as possible during the passive movement phase, which directly preceded our event of interest – location occupation event. Next, the participant’s first-person perspective was smoothly pinned down to vertically face the ground and fixed at the ground for 4 seconds (i.e., location occupation event). Afterwards, they had to report the color of the ball positioned at the current location, by choosing one of the four options displayed on a black blank screen. Importantly, the order of the four options displayed on the screen was randomized from trial to trial, and participants pressed a designated button to cycle through the options. This setup ensured that each test location was not associated with any fixed position on the screen or any consistent pattern of joystick movements across trials. To prevent the use of timing or counting strategies, the movement speed during the passive transport was randomly sampled from a uniform distribution ranging from 2 m/s to 5 m/s on a trial-by-trial basis. Accuracy was emphasized, but participants were instructed to not spend longer time than necessary. Feedback was provided after the participant had made response, telling them whether their response was correct and, if incorrect, what the correct answer was.

This task contrasted landmark-based navigation and path integration. Landmark-based navigation refers to the strategy of relying on the landmark (e.g., the tree) for localization. Path integration refers to the strategy of estimating one’s traveled distance from the fixed starting position (i.e., the red arrow) based on self-motion cues to infer self-location.

The task consisted of three conditions: self-motion condition, landmark condition, and combination condition. To match the sensory inputs across the three cue conditions as much as possible, both the landmark (a tree) and self-motion cues were provided in all cue conditions. In the two single-cue conditions, the position of the tree was adjusted to create spatial conflicts between self-motion and landmark cues, meaning the correct target location defined by the two cues differed. Participants were instructed to use either the landmark information or self-motion information exclusively. Feedback was defined by the task-relevant cue type in the condition to reinforce the designated navigation strategy. Specifically, in the landmark condition, participants were instructed to estimate self-location via their distance to the tree, while ignoring the distance they had traversed from the red arrow. In other words, they were instructed to employ landmark-based navigation exclusively. Conversely, in the self-motion condition, participants were instructed to rely on the traveled distance from the red arrow, while ignoring the tree. In other words, they were instructed to employ path integration exclusively. In the combination condition, the tree was positioned at its original position, meaning no conflict between self-motion and landmark cues, allowing participants to utilize both navigation strategies to identify the test location.

### Experimental procedure

The experiment was conducted over three separate days, with behavioral training on the 1st day (Pre-scan_day) and MRI scanning on the 2nd day (MRI_day1) and 3rd day (MRI_day2). Behavioral training occurred one day before MRI_day1 (Figure 1c). Due to logistic restrictions, the time interval between the two scanning sessions ranged from one to four days.

*<u>Behavioral training (Pre-scan_day)</u>*. The behavioral training aimed to familiarize participants with the VR environment and learn the positions of the four test locations in relation to the landmark (i.e., the tree) and the starting position of path integration (i.e., the red arrow). The training consisted of three parts. Each part consisted of a learning phase, followed by a testing phase. In the learning phase, participants performed the learning task, as described in an earlier paragraph in the Methods section. In the testing phase, participants performed the location identification task. The testing phase had four blocks for each part: one block for landmark and self-motion condition and two for combination condition. For the first part, each testing block included four trials, each trial corresponding to one of the four test locations. For the second part and the third part, each block included 16 trials, with four trials for each test location. Within each testing block, the trials were counterbalanced among the four test locations.

*<u>MRI scanning (MRI_day1 & MRI_day2).</u>* The two MRI scanning sessions followed an identical protocol, starting with a practice session to reacquaint participants with the tasks. The functional scanning phase involved the “Location Identification Task”, with three runs each day. Each run was subdivided into three blocks for each cue condition. The order of conditions was pseudo-randomized, using Latin-square counterbalance with the restriction that two successive runs did not belong to the same cue condition within a day. Functional MRI scans lasted approximately one hour each day, with total scanning time reaching up to 1.5 hours.

Each cue condition included six fMRI blocks of trials, resulting in 6*3 = 18 blocks in total. Each scanning day included nine blocks, organized into three functional runs. We adopted the continuous carry-over design. Using the path-guided cycle, we generated de Brujin sequences, with 2nd order counterbalancing (Aguirre et al., 2011). In the sequence, the order of test locations was balanced, so that each test location had equal probabilities of being preceded by each of the other test locations. Each de Bruijn sequence corresponded to one block of trials. In each de Bruijn sequence, there were five types of events: the location occupation periods at the four test locations, during which participants remained at the test location for 4 seconds, and the null event, where participants fixated their eyes on a cross displayed in the center of the blank screen. Consequently, the de Bruijn sequence comprised 25 events in total, with five repetitions for each event type. To ensure that the hemodynamic response reached a steady state before the sequence commenced, we replicated the final event in the sequence and positioned it at the beginning. Although this duplicated event was modeled in the first-level GLMs, it was excluded from the fMRI adaptation analyses.

We employed the de Brujin cycle generator provided by GK Aguirre (https://www.cfn.upenn.edu/aguirre/wiki/doku.php?id=public:de_bruijn_software).The generator uses path-guided algorithm to find the sequence with high detection power. We selected nine de Brujin sequences with high detection power (> 0.66) and low correlation coefficient (auto correlation < 0.02). The nine original sequences were randomly assigned to the three cue conditions, with three sequences for each cue condition. For each cue condition, we further generated three altered de Bruijn sequences based on the three original ones, resulting in six sequences for each cue condition. For each cue condition, the six sequences were randomly assigned to the six blocks. Consequently, each block consisted of 20 effective trials for the fMRI adaptation analyses, resulting in 20 * 6 (blocks) = 120 trials for each cue condition in total and 120 * 3 (cue conditions) = 360 trials in total in the entire scanning procedure.

In the single-cue conditions, the two cue types indicated different target locations and participants were required to attend to one cue while ignoring the other. For the single-cue conditions, we were primarily interested in the neural representations of spatial locations defined by the task-relevant cue type. Nevertheless, when designing the experiment, we had a secondary interest of examining whether the brain also represents spatial location information defined by the task-irrelevant cue, motivated by the frequent behavioral finding that path integration seems to be operate automatically without voluntary attention (Gallistel, 1990; Shettleworth & Sutton, 2005; but see Chen & Mou, 2024; Zhao & Warren, 2015 for alternative views). Therefore, the nine de Bruijn sequences were transformed and assigned to blocks in such a way that the detection power for fMRI adaptation was matched between the task-relevant cue and the task-irrelevant cue across the six blocks for each single-cue condition. Crucially, the detection power for fMRI adaptation was matched among the three cue conditions (landmark, self-motion, and combination), so that any differences among the cue conditions in adaptation could not be attributed to differences in detection power. The procedure for generating the sequences is depicted in Figure S7.

### MRI acquisition and preprocessing

Imaging data were acquired using a 3T SIEMENS (Erlangen, Germany) Magnetom Prisma scanner, with a 64-channel phased array head coil. Scans comprised a whole-head, three-dimensional structural T1weighted anatomical image with 1 mm isotropic resolution (repetition time (TR)/ echo time (TE)/inversion time = 2300/2.32/900 ms; flip angle = 8°; field of view (FOV) = 240 × 240 mm; 208 slices; GRAPPA acceleration factor 1); a high resolution moderately T2-weighted structural image comprising the hippocampus and EC acquired perpendicular to the long axis of the hippocampus using a turbospin-echo sequence (in-plane resolution = 0.4 × 0.4 mm, slice-thickness = 1.5 mm; TR/TE = 4700 ms/52 ms; FOV = 224 × 224 mm; 34 slices); six runs of T2*-weighted functional images acquired with a partial volume echo-planar imaging sequence, aligned with the long axis of the hippocampus (in-plane resolution = 1.5 × 1.5 mm, slice-thickness = 1.5 mm; TR/TE = 2000/30 ms; flip angle = 90°; FOV = 192 × 192 mm; 27 slices; GRAPPA acceleration factor 1); 10 volumes of whole-brain functional scans (TR/TE = 8200 ms/30ms; flip angle = 90°; FOV = 192 × 192 mm; 118 slices; resolution = 1.5 mm isotropic), which were used to facilitate the co-registration of anatomical masks segmented on the T2-weighted structural scan to the functional scans.

The T1-weighted structural images were subjected to bias correction using Advanced Normalization Tools (ANTs). The functional images are initially realigned by matching the first scan from each session to the first scan of the first session. Subsequently, the images within each session are aligned to the session’s first image. The functional images were left unsmoothed at this stage. In the later ROI-based analyses, beta estimates for effects of interest, such as location-based adaptation, were averaged across all voxels within each ROI, yielding an aggregate effect estimate for the entire region. For voxel-wise analyses, beta images were first spatially smoothed with a 3 mm isotropic full-width-at-half-maximum (FWHM) to reduce the impact of noise and alleviate anatomical variability across participants, before being normalized to the Montreal Neurological Institute (MNI) template.

### Anatomical masks for regions of interest

In this study, we focused on two primary areas of the brain as our regions of interest: the retrosplenial cortex (RSC) and various regions within the medial temporal lobe (MTL), including the hippocampus, parahippocampal cortex (PHC), entorhinal cortex (EC), and perirhinal cortex (PRC). The anatomical masks for these areas in an example participant’s brain are displayed in Figure 3a. These areas are critical for memory and navigation, making them of particular interest for our investigation.

The RSC mask was automatically extracted from each participant’s T1-weighted structural scan using the Freesurfer software. RSC was defined as the posterior-ventral portion of the cingulate gyrus, which mainly consists of BA29/30. Note that the definition of RSC is anatomically different from the retrosplenial complex, which is a functionally defined region typically extending into the parieto-occipital sulcus (Epstein, 2008). The MTL regions, including the hippocampus, PHC, EC, and PRC, were segmented manually on each participant’s high-resolution T2-weighted structural scan (in-plane resolution = 0.4 × 0.4 mm) in ITK-SNAP (version 4.0 http://www.itksnap.org/pmwiki/pmwiki.php). This manual segmentation followed a detailed protocol developed by Berron, Vieweg, and colleagues (Berron et al., 2017). EC was further divided into anterior-lateral EC and posterior-medial EC, following the procedure used in our previous study (Chen et al., 2019).

The hippocampus was further divided into the anterior and posterior portions on the T2-weighted structural scan in ITK-SNAP. The two portions comprised an equal number of coronal slices of the hippocampus. When the total number of coronal slices was odd, the anterior portion contained one more slice than the posterior portion.

For completeness, the hippocampus was also divided into six subfields in ITK-SNAP: CA1, CA2, CA3, subiculum (SUB), dentate gyrus (DG), and the hippocampal tail, also following the procedure developed by Berron, Vieweg, and colleagues (Berron et al., 2017). In functional data, CA2 could not be reliably separated delineated, due to its small volume. Therefore, CA2 and CA3 were combined (CA2/CA3), in accordance with standard practices in previous 3T fMRI studies with similar scanning protocols (Bonnici et al., 2012; Chadwick et al., 2014). Results of location-based adaptation for hippocampal subfields are illustrated in Figure S5.

The anatomical masks for the MTL regions were co-registered to the mean functional scan of the first scanning day in SPM12, using the following procedure: first, the mean whole-brain functional scan was co-registered to the mean functional scan; second, the T2-weighted structural scan, along with the anatomical masks, were co-registered to the mean whole-brain functional scan obtained from the first step; third, the co-registered anatomical masks were re-sliced using nearest-neighbor interpolation, with the mean functional scan as the reference image. The anatomical mask for RSC, along with the T1-weighted structural scan, was first co-registered to the mean whole-brain functional scan, which had already been co-registered to the mean functional scan; then the co-registered anatomical mask was resliced using nearest-neighbor interpolation, with the mean functional scan as the reference image.

### Behavioral analyses

Behavioral data analysis focused on participants’ accuracy in identifying the test locations. Correct responses were coded as 1, and incorrect responses as 0. Given four color options in the task design, the chance level of behavioral accuracy was 0.25. Behavioral data analyses were conducted using JASP software (JASP Team (2023). JASP (Version 0.17.1)).

### Cognitive modeling: Extended signal detection model

To dissociate representational precision from response bias and attentional failure in behavioral performance, we applied an extension of signal detection theory to our location identification task with four choices. The modeling procedure was similar to the one adopted in our previous study, but with a few improvements. The three cue conditions were fit separately.

In each cue condition, there were four test locations *Loc_i_*(*i* ∈ [1, 2, 3, 4]). We included eight free parameters to model the four standard deviations of the underlying representations of the four test locations σ_*i*_ (*i* ∈ [1, 2, 3, 4]), the three response criteria *C*_*i*,(*i*+1)_ (*i* ∈ [1, 2, 3]), and the lapse rate (*lr*). The response criterion *C*_*i*,(*i*+1)_ represents the choice boundary between *Loc*_*i*_ and *Loc*_*i*+1_. The lapse rate represents the proportion of trials in which participants completely failed in attention and simply chose a response randomly (Zhang & Luck, 2008). The centers of the representation distributions μ*i* (*i* ∈ [1, 2, 3, 4]) were assumed to be at the true positions of the test locations (i.e., *μ*_1_= -3 m, *μ*_2_*=* -1 m, *μ*_3_*=* 1 m, and *μ*_4_*=* 3 m).

In each round of simulation, given each set of althorigm-generated values of the free parameters, we calculated the 4 X 4 predicted behavioral confusion matrix, with each element denoting the probability of each location-response pair. This matrix can be solved analytically. Given a test location *Loc*_*i*_ physically occupied by the participant, the probabilities the participant judges it to be each of the four test locations are calculated based on the cumulative normal distributions of the physical location and the response criteria. Considering the lapse rate, the probability of juding *Loc*_*i*_ as *Loc*_1_ is:

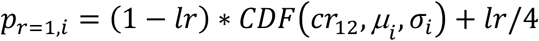

where CDF stands for the cumulative distribution function of the normal distribution, *cr*_12_ denotes the threshold to calculate the area of the cumulative normal distribution below *cr*_12_, μ_*i*_ and σ_*i*_ represent the mean and standard deviation of the cumulative normal distribution.

The probability of juding *Loc*_*i*_ as *Loc*_2_ is:

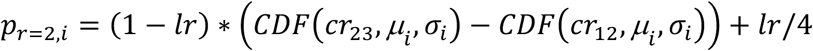

The probability of juding *Loc*_*i*_ as *Loc*_3_ is:

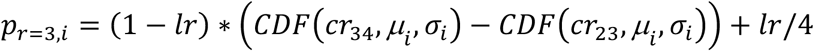

The probability of juding *Loc*_*i*_ as *Loc*_4_ is:

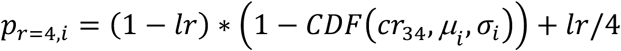

We then compared the actual behavioral confusion matrix to the predicted confusion matrix, by computing the probability of observing each actual response given the predicted confusion matrix. Finally, we summed the log-transformed probabilities of all actual responses,

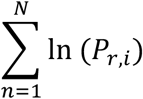

in which n represents the trial number and N represents the total number of trials in the actual experiment. If *P*_*r,i*_ equaled to 0, we manually set it to be extremely small (=10^-12^), otherwise ln (*P*_*r,i*_) would become illegitimate.

Note that in our previous application of this model (Chen et al., 2024), we obtained the predicted behavioral confusion matrix by simulating a large number of trials (=1000 trials for each test location). This method is valid but time-consuming.

We repeated the simulation to maximize the log-transformed probability (i.e., maximum likelihood estimation) summed across four test locations. We used the Hooke & Jeeves hill-climbing algorithm for model optimization (Hooke & Jeeves, 1961), as implemented in Matlab_R2020a. To avoid the potential local-minima problem, the model-fitting procedure was repeated 20 times with randomized starting values for the parameters each time, and the parameter estimates with the best fit were selected.

### Cognitive modeling: Comparing cue-handling strategies in combination condition

Four different models were compared, corresponding to four cue-handling strategies: Bayesian cue integration, Bayesian cue alternation, landmark dominance, and self-motion dominance. We constructed two different sets of models for these four strategies, with one set assuming sensory biases and the other assuming no sensory biases.

When sensory biases were assumed, each model consists of 28 free parameters: sensory standard deviations for the four test locations in the landmark condition (σ_1,*land*_, σ_2,*land*_, σ_3,*land*_, σ_4,*land*_), sensory standard deviations for the four test locations in the self-motion condition (σ_1,*motion*_, σ_2,*motion*_, σ_3,*motion*_, σ_4,*motion*_), sensory biases for the four test locations in the landmark condition (*μ*_1,*land*_, *μ*_2,*land*_, *μ*_3,*land*_, *μ*_4,*land*_), sensory biases for the four test locations in the self-motion condition (*μ*_1,*motion*_, *μ*_2,*motion*_, *μ*_3,*motion*_, *μ*_4,*motion*_), response criteria in the landmark condition (*Cr*_12,*land*_, *Cr*_23,*land*_, *Cr*_34,*land*_), response criteria in the self-motion condition (*Cr*_12,*motion*_, *Cr*_23,*motion*_, *Cr*_34,*motion*_), response criteria in the combination condition (*Cr*_12,*comb*_,*Cr*_23,*comb*_, *Cr*_34,*comb*_), attentional lapse rate in the three cue conditions (*lr*_*land*_, *lr*_*motion*_, *lr*_*comb*_).

When sensory biases are not assumed, the centroids of the sensory measurement distributions for the four test locations in both single-cue conditions are fixed at the target locations (i.e., *μ*_1_= -3 m, *μ*_2_*=* -1 m, *μ*_3_*=* 1 m, and *μ*_4_*=* 3 m), resulting in 20 free parameters in total for each model.

In the Bayesian cue integration model, for each test location, the weights assigned to spatial cues are determined by cue relatively reliability:

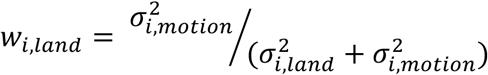

The sensory bias for location *i* in the combination condition is calculated as:

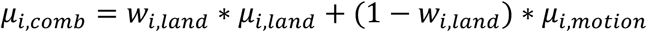

The sensory standard deviation for location *i* in the combination is calculated as:

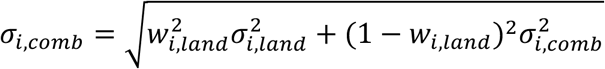

In the Bayesian cue alternation model, weights assigned to the two cue types are the same as in the Bayesian cue integration model, so is the calculation of sensory biases in the combination condition. The key difference lies in the calculation of sensory standard deviations in the combination condition. The sensory standard deviation for location *i* in the combination is calculated as:

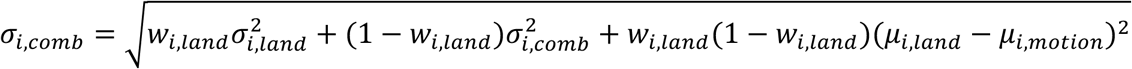

Since weights are smaller than 1, the predicted sensory standard deviations in the combination condition are larger for the Bayesian cue alternation model than for the Bayesian cue integration model, even when there exists no disparity in sensory bias between the two cue types at each test location (i.e., *μ*_*i*,*land*_ − *μ*_*i*,*motion*_ = 0). This difference reflects the benefit of precision enhancement brought by cue integration.

In the landmark dominance model, sensory biases and sensory standard deviations in the combination condition are the same as those in the landmark condition:

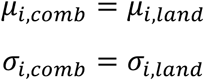

Similarly, in the self-motion dominance model, sensory biases and sensory standard deviations in the combination condition are the same as those in the self-motion condition:

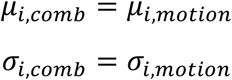

The model fitting procedure was the same as that in the modeling analysis that fit the extended signal detection model to the three cue conditions separately (see Methods - Cognitive modeling: Extended signal detection model). The model fitting procedure maximized the log-transformed probability (i.e., maximum likelihood estimation) summed across four test locations and three cue conditions.

### fMRI analysis of location-based adaptation

*<u>First-level analysis.</u>* To assess fMRI adaptation in relation to inter-location distance, we constructed a separate first-level GLM (*GLM-adaptation-location*). Along with the event regressors that modeled the location occupation phase (Figure 1b), we included parametric regressors modeling the modulatory effects of spatial distance between sequentially visited locations. These parametric regressors captured the continuous variation of inter-location distance, specifically with values of 0, 4, 8, and 12 m. Separate regressors were created to model the location occupation periods that were not suitable for adaptation, that is, being preceded by a null event or being in the first trial. Same as in *GLM-succes*s, to control for potential effects of the passive movement stage on the adaptation effects associated with the ensuing location occupation period, we included regressors modeling the passive movement phase, irrespective of the test location. Head motion parameters (three rotation parameters and three translation parameters) were included as nuisance regressors. Volumes in which head motion exceeding the threshold defined by the Carling’s box-plot rule were modeled with separate spike regressors (Siegel et al., 2014). Event regressors were convolved with the canonical hemodynamic response function (plus temporal derivatives). Scans were not concatenated across runs, and a constant term was included to model baseline activity for each run.

To visualize the adaptation effects, we constructed a separate first-level GLM (*GLM-adaptation-visualization*), in which separate regressors modeled the location occupation periods with different inter-location distances (i.e., 0, 4, 8, and 12 m). The beta estimates of these regressors were then plotted as a function of inter-location distance.

*<u>Second-level ROI-based analysis.</u>* First, beta estimates of adaptation were averaged across all voxels within each ROI. Next, directional one-sample t-tests were conducted on the participant-specific beta estimates of adaptation for each cue condition (landmark, self-motion, combination). We performed one-sided statistical tests because we hypothesized that BOLD responses should decrease as the distance between successively visited locations decreased, that is, positive adaptation. In addition, negative adaptation is difficult to interpret (Barron et al., 2016; but see Garvert et al. (2023)). The nonparametric permutation-based maximum t statistic approach was adopted to correct for multiple comparisons.

For each group-level t test, we calculated the Bayes factor (BF10), which indicates the relative likelihood of the alternative hypothesis (i.e., the group mean was > 0) over the null hypothesis (i.e., the group mean was not > 0; Rouder et al., 2009). The effect size scale (r) adopted was 0.707. A BF_10_ greater than 3/10/30 indicates moderate/strong/very-strong evidence for the alternative hypothesis, whereas a BF_10_ less than 0.333/0.1/0.033 indicates moderate/strong/very-strong evidence for the null hypothesis (Jeffreys, 1961). BF_10_ was also calculated in other ROI-based group-level fMRI analyses. Statistical outliers were identified using the boxplot rule, that is, a value would be considered as an outlier if it is larger than 3^rd^ quartile + 3*interquartile range or smaller than 1^st^ quartile − 3*interquartile range. Statistical outliers were winsorized to the nearest inlier (Reifman and Keyton, 2010).

*<u>Second-level voxel-wise analysis.</u>* Voxel-wise analysis is an important supplement to the ROI analysis. Participant-specific adaptation maps were normalized to the Montreal Neurological Institute (MNI) template and spatially smoothed with 3 mm isotropic FWHM. We conducted directional one-sample t tests against 0. Multiple comparisons were corrected using the nonparametric permutation-based approach (Nichols and Holmes, 2002), using the voxel-level inference approach or the cluster-level inference approach (voxel-wise T >3). To explore beyond our ROIs, we corrected for multiple comparisons across the entire volume.

### fMRI analysis of task-based connectivity

*<u>Graph analysis.</u>* First, we assessed the functional connectivity between these regions using the beta-series connectivity analysis. We constructed a first-level GLM (*GLM-connectivity*) to extract the beta-series of brain activation for the location occupation period for each cue condition (Cisler et al., 2014). This GLM used separate event regressors to model the location occupation period of individual single trials. No parametric regressors were included. Other aspects of the GLM were the same as the GLM used to assess the location-based adaptation (i.e., *GLM-adaptation-location*). For each brain region, we obtained a temporal sequence of activation estimates concatenated across individual trials, which were mean-centered within each run prior to the trial concatenation. We then calculated pairwise Pearson r correlations (fisher-transformed) between the temporal sequences of these regions for each participant.

Next, we conducted the graph analysis to explore the structure of the network comprising our pre-defined ROIs (i.e., the MTL regions and RSC), using GRETNA (Wang et al., 2015). To maximize the number of regions, which is critical to ensure a successful graph analysis, we extracted beta time-series for the two hemispheres separately for each ROI. In addition, alEC and pmEC were analyzed separately, resulting in a total number of 6*2 = 12 regions. A network consists of nodes and edges: nodes represent individual brain regions, and edges represent the strength of functional connectivity between nodes. Inter-region functional connectivity was calculated based on beta-series of brain activation (Cisler et al., 2014).

We calculated group-level functional connectivity matrix for each cue condition. The matrices were then thresholded at 60%, meaning 60% of edges were retained, as this is the minimum proportion to ensure that every node had at least one inter-node connection. We used the same thresholding method when analyzing participant-specific functional connectivity matrices.

We adopted a greedy optimization algorithm to find the network partition with the highest modularity value (Chen et al., 2008; Danon et al., 2006). Hubs are nodes that play critical roles in connecting different parts of the network. To identify hubs, we measured betweenness centrality for each node, which equaled to the number of shortest paths passing through the node. A node was identified as a hub if its betweenness centrality exceeding one standard deviation above the mean averaged across all nodes in the network (Kong et al., 2017). We were particularly interested in nodal hubness, because a hub receives various information sources from various regions, likely rendering it the locus of cue integration (van den Heuvel & Sporns, 2013).

*<u>Generalized psychophysiological</u>* <u>interaction (gPPI) *analysis*</u>. The aforementioned network analysis is based on mean time series of activation for each ROI. It is possible that potential effects were relatively restricted in space, which would be washed away in the ROI-based analysis. To overcome this potential limitation, we conducted the gPPI analysis, which is suitable to revealing context-modulated functional connectivity effects at the voxel-level (Cisler et al., 2014). We performed the gPPI analysis, which was developed to contrast connectivity strength between more than two conditions, as was the case in the current study. Since the fMRI adaptation analysis showed that the hippocampus showed strong and robust spatial coding in the combination condition (see Results), the hippocampus was used as the seed region.

In the gPPI analysis, first, the BOLD signals of the seed region as well as the stimulus conditions (i.e., the three cue conditions) were deconvolved to obtain the neural event estimates for the hippocampus (*x*_*hipp*_) and the three cue conditions (*x*_*land*_, *x*_*motion*_, *x*_*comb*_). Next, interaction terms were created between the neural event estimates of the hippocampus and each individual cue condition (i.e., *x*_*hipp*_ \**x*_*land*_, *x*_*hipp*_ \**x*_*motion*_, *x*_*hipp*_ \**x*_*comb*_). Finally, we calculated the differences between the combination condition and the two single-cue conditions, by subtracting the corresponding interaction terms. For example, to investigate whether any voxels displayed stronger connectivity with the hippocampus in the combination condition than the landmark condition, we tested the statistical significance of the term *x*_*hipp*_\**x*_*comb*_ - *x*_*hipp*_\**x*_*land*_.

Same as the abovementioned voxel-based analyses of the fMRI adaptation analysis, participant-specific result maps were normalized to the MNI template and spatially smoothed with 3 mm isotropic FWHM. Because we expected stronger connectivity strength with the hippocampus in the combination condition than the two single-cue conditions (Cooper & Ritchey, 2020; Jakobs et al., 2012; Watanabe et al., 2001), we conducted directional one-sample t tests against 0. Multiple comparisons were corrected using the nonparametric permutation-based approach (Nichols and Holmes, 2002), using the voxel-level or cluster-level inference approach (voxel-wise T > 3). To examine voxels within our pre-defined ROIs, we used group-level masks comprising of our pre-defined ROIs for small volume correction. To explore beyond our ROIs, we corrected for multiple comparisons across the entire volume, also using the nonparametric permutation-based test.

### Adaptation-based neural space reconstruction analysis

To evaluate whether the fMRI adaptation effect contains fine-grained distance information between different test locations, we adapted the neural space reconstruction analysis to fMRI adaptation, the same method employed in our previous report (Chen et al., 2024).

This analysis consisted of three steps. In Step 1, we constructed the neural distance matrix, whose elements denote pairwise neural distances between the test locations. The neural distance between two test locations could be quantified as the brain region’s activation level for one location when preceded by the other location - the lower the region’s activation to the current location when preceded by the other location, the larger the repetition suppression effect, the closer the two test locations would be positioned to each other in the neural space. We constructed a GLM, which modeled the location occupation phase in individual single trials with separate regressors. To construct the neural distance matrix, these trials were classified into 4×4 = 16 groups based on the combination of two locations visited in succession; the beta estimates for trials from the same group were averaged, resulting in the 4×4 neural distance matrix. In the matrix, rows represent the previous location, columns represent the current location, and each element represents the activation level at the current location when preceded by the previous location.

To keep it consistent with the main fMRI adaptation analysis, the neural space reconstruction analysis was restricted to trials that could be modeled for adaptation (i.e., locations not preceded by the null event and not the first event in the sequence).

In Step 2, we normalized the neural distance matrix to render all the elements within the range [0, 1]. Elements that were diagonally symmetrical to each other in the matrix were averaged.

In Step 3, the normalized neural distance matrix was subjected to multidimensional scaling and the Procrustes analysis. The multi-dimensional scaling recovers the spatial coordinates of the locations in the neural space, following the basic principle that locations with greater representational similarity are positioned closer to each other in the neural space (Kruskal & Wish, 1978). Subsequently, the Procrustes analysis mapped the estimated coordinates of the locations to the original physical space through rotations and reflections (Gower & Dijksterhuis, 2004).

To assess whether the neural space resembled the original physical space, we adopted a nonparametric permutation-based test. The normalized neural distance matrix was averaged across participants to obtain the grand group-level neural distance matrix for the four test locations (Marchette et al., 2014; Peer & Epstein, 2021; Persichetti & Dilks, 2019), which was then subjected to multidimensional scaling and the Procrustes analysis. The nonparametric permutation test was conducted as follows. First, we obtained the actual Procrustes distance calculated from the group-level neural distance matrix, which indicates the deviation of the reconstructed neural space from the original physical space. Second, we applied the permutation procedure to obtain the surrogate distribution of Procrustes distance, to which the actual Procrustes distance would be compared. Specifically, in each permutation, we randomly shuffled the entries in the grand group-level neural distance matrix. Note that to allow for more permutations, this shuffling was done prior to the averaging of symmetrical off-diagonal elements in the neural distance matrix. We obtained the Procrustes distance by applying multidimensional scaling and the Procrustes analysis to the shuffled neural distance matrix. This process was repeated 5000 times, resulting in a surrogate distribution of Procrustes distance. Third, the actual Procrustes distance was compared to the surrogate distribution. The significance level (i.e., p value) was calculated as the proportion of values in the surrogate distribution being smaller than the actual Procrustes distance, analogous to directional one-sample t test. Significant results would indicate that the group-level neural space resembled the original physical space.

### Analysis of empirical relative detection power of fMRI adaptation

In the current study, the de Bruijn sequences were generated based on true locations, and hence, were theoretically optimized in terms of *DP_rel_* with respect to objective locations. However, participants’ responses or trial-wise path lengths could not be known in advance, leading to the possibility that *DP_rel_* was reduced for the response-defined sequences compared to the location-defined sequences.

To address this issue, we calculated the empirical *DP_rel_* for location-defined and response-defined sequences separately, based on the first-level general linear models (GLMs). These GLMs included regular regressors modeling the location occupation events as a measure of the direct stimulus effect, and a parametric regressor modeling objective location, or subjective response as a measure of the adaptation effect.

Specifically, *DP_rel_* is calculated as follows (Aguirre et al., 2011),

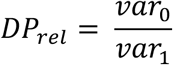

in which *var*_0_ represents the hypothesized neural modulation after HRF convolution and high-pass filtering (e.g., f > 1/128), and *var*_1_ represents the original hypothesized neural modulation prior to HRF convolution and high-pass filtering.

To calculate *var*_1_, we constructed these first-level GLMs with no HRF convolution and no high-pass filtering applied. We then converted the simulated BOLD signal of the parametric regressor from the time domain to the frequency domain, using the fast Fourier transform (FFT). The variance of the hypothesized neural modulation for the parametric regressor (i.e., *var*_1_) was calculated as the area under curve (AUC) using the frequency-domain data.

To calculate *var*_0_, we convolved the predicted fMRI time series for the parametric regressor with the canonical hemodynamic response function (HRF), and adopted a high-pass filter with a cut-off at 1/128s = 0.0078 Hz. The variance of the convolved and filtered signal for the parametric regressor (i.e., *var*_0_) was calculated in the same way as *var*_1_. Finally, to obtain *Dp_rel_*, we divided *var*_0_ over *var*_1_.

## Supplemental Information

**Figure S1.**
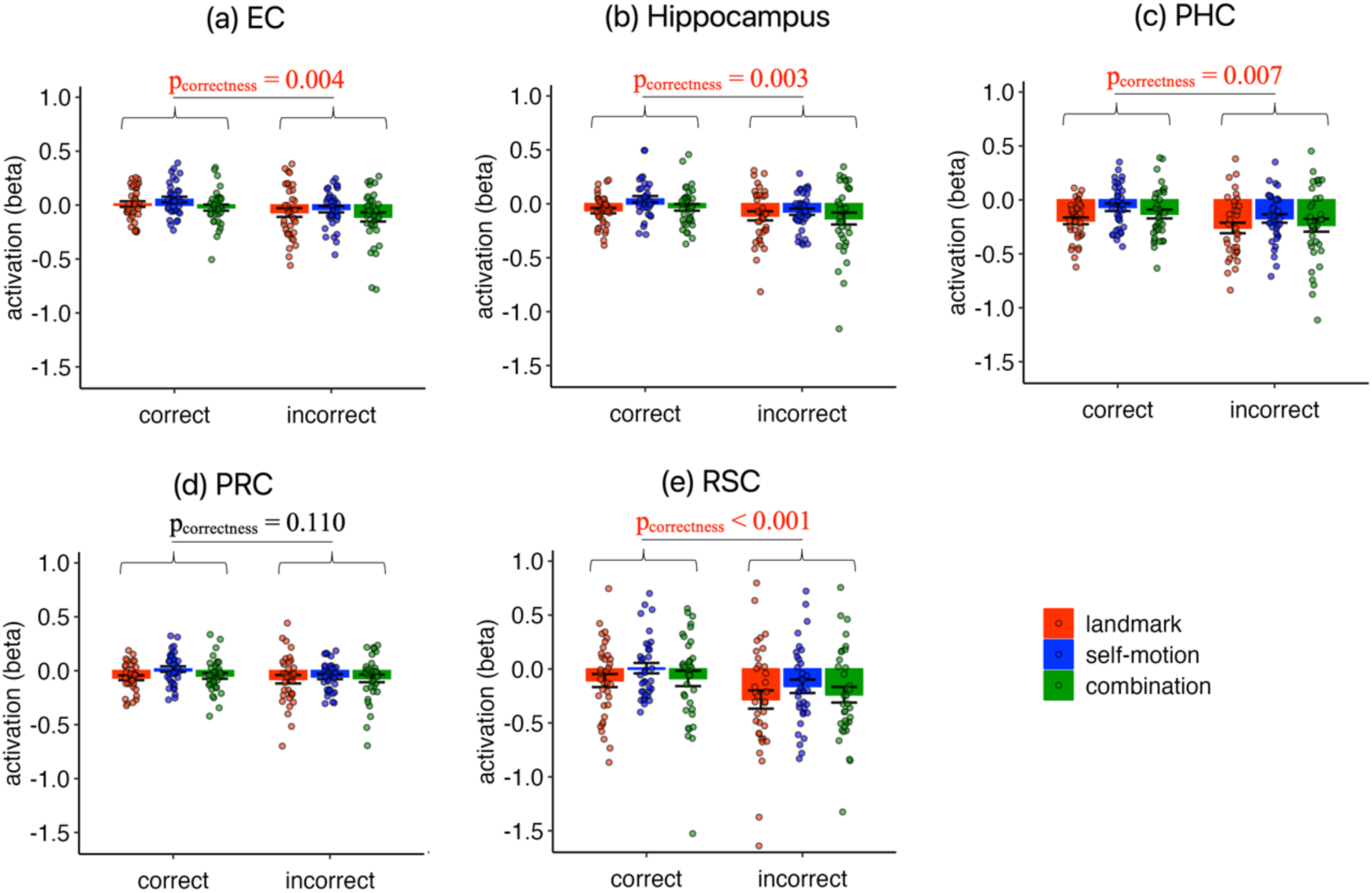
Effects of successful navigation in all ROIs. Brain activation is plotted as a function of cue condition (landmark vs. self-motion vs. combination) and navigation success (correct vs. incorrect). Brain activation corresponds to the beta weight estimated for the event regressor modeling the location occupation period in the location identification task. The hippocampus, EC, RSC, and PHC showed significant main effect of navigation success. Correct trials showed stronger activation than incorrect trials in the hippocampus (main effect of correctness, F(1,34) = 10.555; p = 0.003, 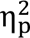 = 0.237), EC (F(1,34) = 9.586; p = 0.004, 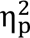 = 0.220), PHC (F(1,34) = 8.234; p = 0.007, 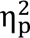 = 0.195), and RSC (F(1,34) = 15.155; p < 0.001, 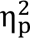 = 0.308), but not in PRC (F(1,34) = 2.706; p = 0.109, 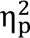 = 0.074; one outlier winsorized, F(1,34) = 2.622; p = 0.115, 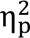 = 0.072). Statistical outliers were highlighted in dark circles, which were winsorized to the nearest inlier in statistical tests. Error bars represent ±SE.

**Figure S2.**
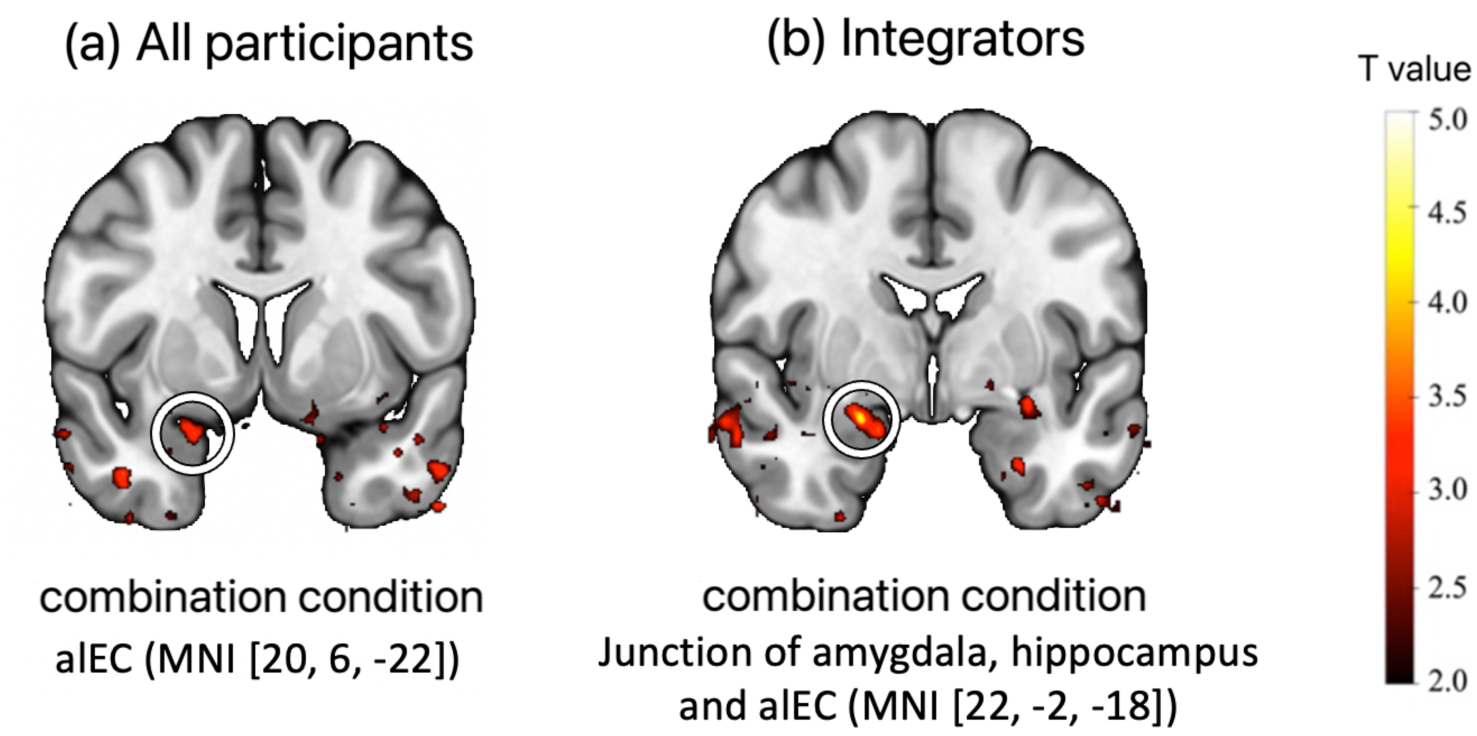
Voxel-wise analysis of location-based adaptation within EC. Participant-specific maps of location-based adaptation were normalized to the MNI template and spatially smoothed with 3mm isotropic FWHM. The parametric t maps were overlaid on the MNI template and projected to the brain surface. We conducted the non-parametric permutation-based test, with the cluster-level inference approach (cluster-defining threshold: T > 3). We used the group-level bilateral EC anatomical mask for small volume correction, for a prior hypothesis that alEC might participate in cue integration (Doan et al., 2019). **(a)** We detected a significant cluster in the combination condition when analyzing all participants (peak T = 3.78, MNI [20, 6, -22], p_FWE,corr,1-tailed_ = 0.048). **(b)** We detected a significant cluster in the combination condition when analyzing integrators only (peak T = 5.18, MNI [22, -2, -18], p_FWE,corr,1-tailed_ = 0.010). For illustrative purposes, the results are displayed on the MNI template (thresholded at T > 2), accompanied by the MNI coordinates of the peak voxels.

**Figure S3.**
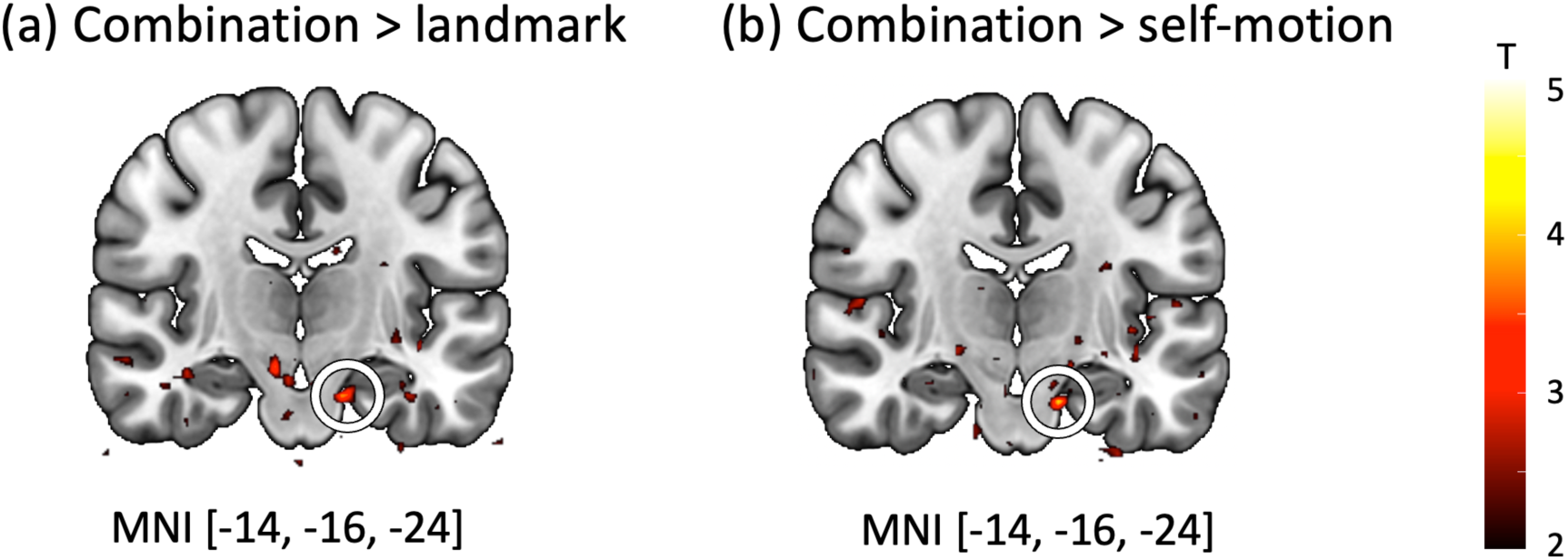
Results of gPPI analysis within EC, using hippocampus as seed region. We conducted the non-parametric permutation-based test. We used the group-level bilateral EC anatomical mask for small volume correction. We detected significantly stronger connectivity with the hippocampus in the combination condition compared to both the landmark condition (MNI [-14, -16, -24], T = 4.34, p_FWE,corr,1-tailed_ = 0.049) and the self-motion condition (MNI [-14, -16, -24], T = 4.57, p_FWE,corr,1-tailed_ = 0.02). Results are overlaid on the MNI template. For illustrative purposes, results are thresholded at T > 2, accompanied by the MNI coordinates of the significant voxels.

**Figure S4.**
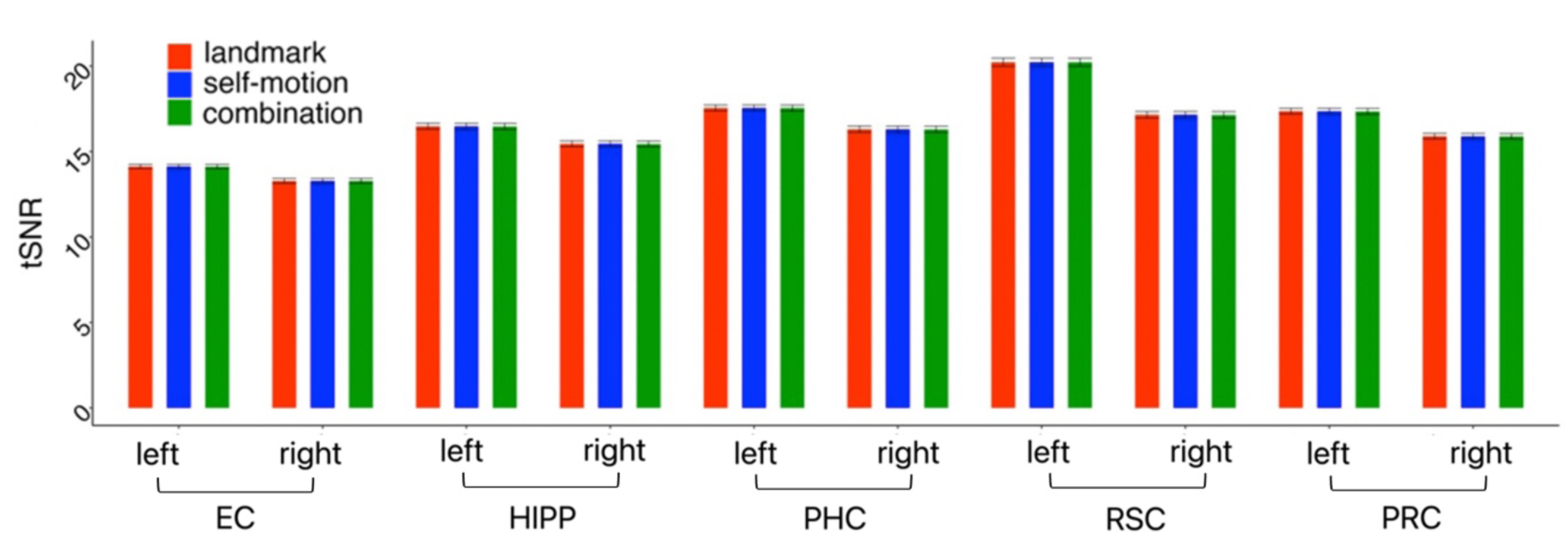
Temporal signal-to-noise ratio (tSNR) in ROIs. Temporal signal-to-noise ratio (tSNR) was calculated for each voxel, which was then averaged across all voxels in the brain region.We submitted tSNR to a repeated-measures ANOVA test, with brain region, hemisphere, and cue type as independent variables. A significant interaction effect was found between brain region and hemisphere (F(4, 136) = 55.365, p < 0.001, 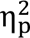 = 0.62). Follow-up analyses showed that the tSNR in the left hemisphere was consistently higher than in the right hemisphere across all ROIs (RSC: F(1, 34) = 470, p < 0.001; EC: F(1, 34) = 80.366, p < 0.001; hippocampus: F(1, 34) = 48.11, p < ; PHC: F(1, 34) = 62.56, p < 0.001; PRC: F(1, 34) = 78.255, p < 0.001). Besides, both the main effects of brain region and hemisphere were significant (brain region: F(4, 136) = 425.42, p < 0.001, 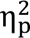 = 0.926; hemisphere: F(1, 34) = 261.413, p < 0.001, 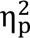 = 0.885). Post-hoc comparisons revealed that tSNR in RSC was higher than other regions (RSC vs EC: t(34) = 40.288, p < 0.001; RSC vs HIPP: t(34) = 22.024, p < 0.001; RSC vs PHC: t(34) = 14.235, p < 0.001; RSC vs PRC: t(34) = 16.671, p < 0.001), tSNR in PHC was higher than EC, PRC and the hippocampus (EC: t(34) = 26.054, p < 0.001; PHC: t(34) = 2.436, p = 0.016; HIPP: t(34) = 7.789, p < 0.001), tSNR in PRC was higher than EC (t(34) = 23.617, p < 0.001) and hippocampus (t(34) = 5.353, p < 0.001) and the tSNR in hippocampus was higher than EC (t(34) = 18.264, p < 0.001). The main effect of cue type was not significant (F(2, 68) = 0.877, p = 0.421, 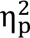 = 0.025). Error bars represent ±SE.

**Figure S5.**
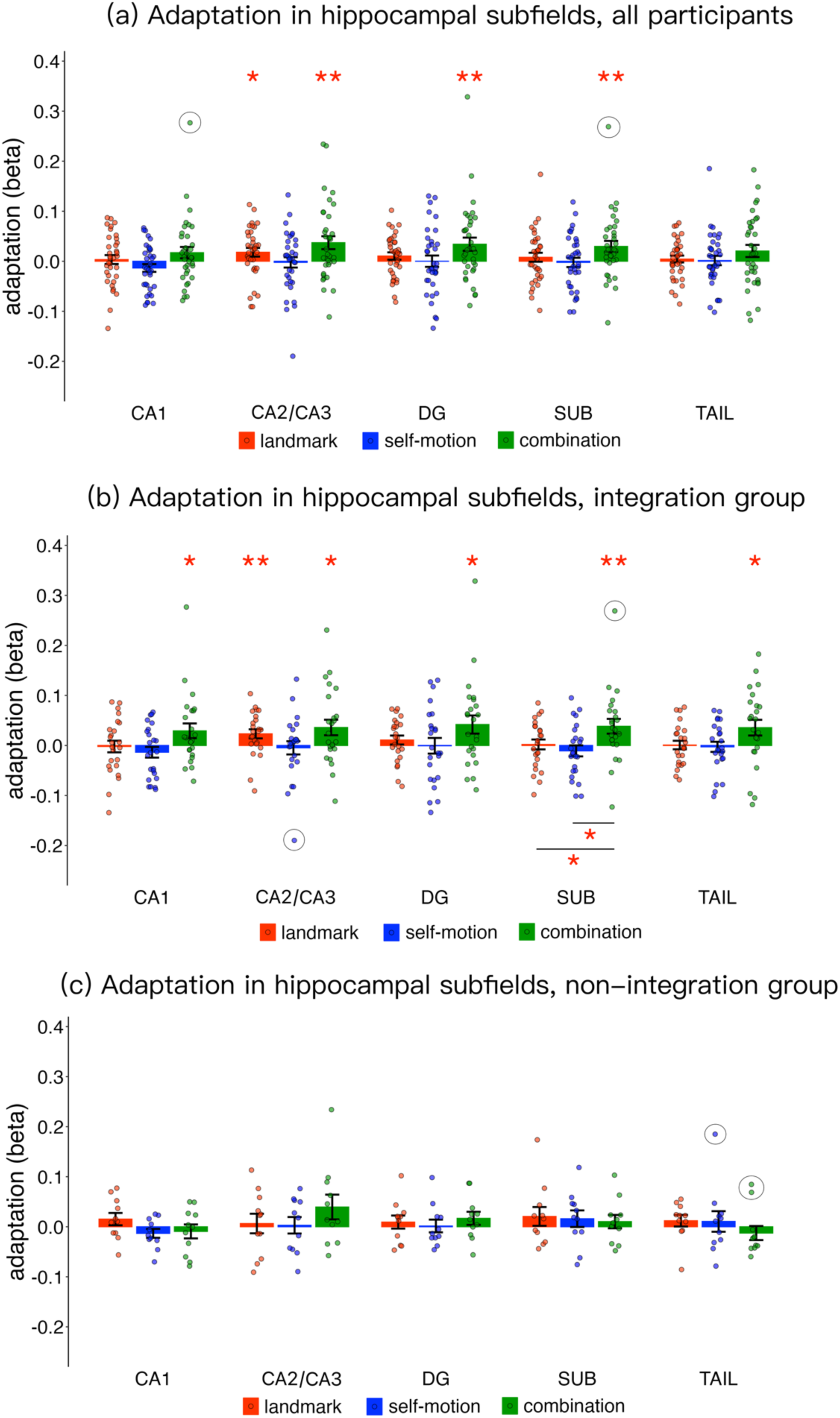
Location-based adaptation in hippocampal subfields. **(a)** Results for all participants. Significant adaptation effects in the combination condition were detected in CA2/CA3 (t(34) = 2.863, p_1-tailed_ = 0.004, BF_10_ = 11.343), DG (t(34) = 2.589, p_1-tailed_ = 0.007, BF_10_ = 6.347), and SUB (t(34) = 2.672, p_1-tailed_ = 0.006, BF_10_ = 7.537). These effects remained significant after multiple comparisons correction across five subfields (ps_corrected_ < 0.04). However, since CA2/CA3 also showed significant adaptation effect in the landmark condition(t(34) = 2.064, p_1-tailed_ = 0.023, BF_10_ = 2.317), which did not differ from the combination condition ((34) = - 1.413, p_1-tailed_ = 0.083, BF_10_ = 0.818), its role in cue integration remains ambiguous. **(b)** Results for integrators. All subfields showed significant adaptation effects in the combination condition : CA1, t(23) = 1.975, p_1-tailed_ = 0.030, BF_10_ = 2.164; DG, t(23) = 2.307, p_1-tailed_ = 0.015, BF_10_ = 3.796; SUB, t(23) = 2.606, p_1-tailed_ = 0.008, BF_10_ = 6.544; Tail, t(23) = 2.271, p_1-tailed_ = 0.016, BF_10_ = 3.563; CA2/CA3 (t(23) = 2.324, p_1-tailed_ = 0.015, BF_10_ = 3.913). After multiple comparisons correction across five subfields, only adaptation in SUB remained significant (p_1-tailed,corrected_ = 0.040). Moreover, adaptation in SUB was significantly greater in the combination condition than in either single-cue condition (vs. landmark, t(23) = 1.998, p_1-tailed_ = 0.029, Cohen’s d = 0.408, BF_10_ = 2.246; vs. self-motion, t(23) = 2.322, p_1-tailed_ = 0.015, BF_10_ = 3.900), reinforcing its role in cue integration. As with the analysis on all participants, CA2/CA3 also showed adaptation in the landmark condition (t(34) = 2.550, p_1-tailed_ = 0.009, BF_10_ = 5.898), which did not differ from the combination condition (t(34) = 0.713, p_1-tailed_ = 0.241, BF_10_ = 0.405), leaving its specific contribution to cue integration unclear. **(c)** Results for non-integrators, no significant adaptation effects were detected in any subfields across cue conditions (ts < 1.6, ps_1-tailed_ > 0.07, BFs_10_ < 1.5). Location-based adaptation corresponds to the beta weight estimated for the parametric regressor modeling the modulatory effect of inter-location distance on BOLD response. Dots represent data from individual participants. Statistical outliers were highlighted in black circles, which were winsorized to the nearest inlier in statistical tests. “*” - p_1-tailed_ < 0.05, “**” - p_1-tailed_ < 0.01. Error bars represent ±SE.

**Figure S6.**
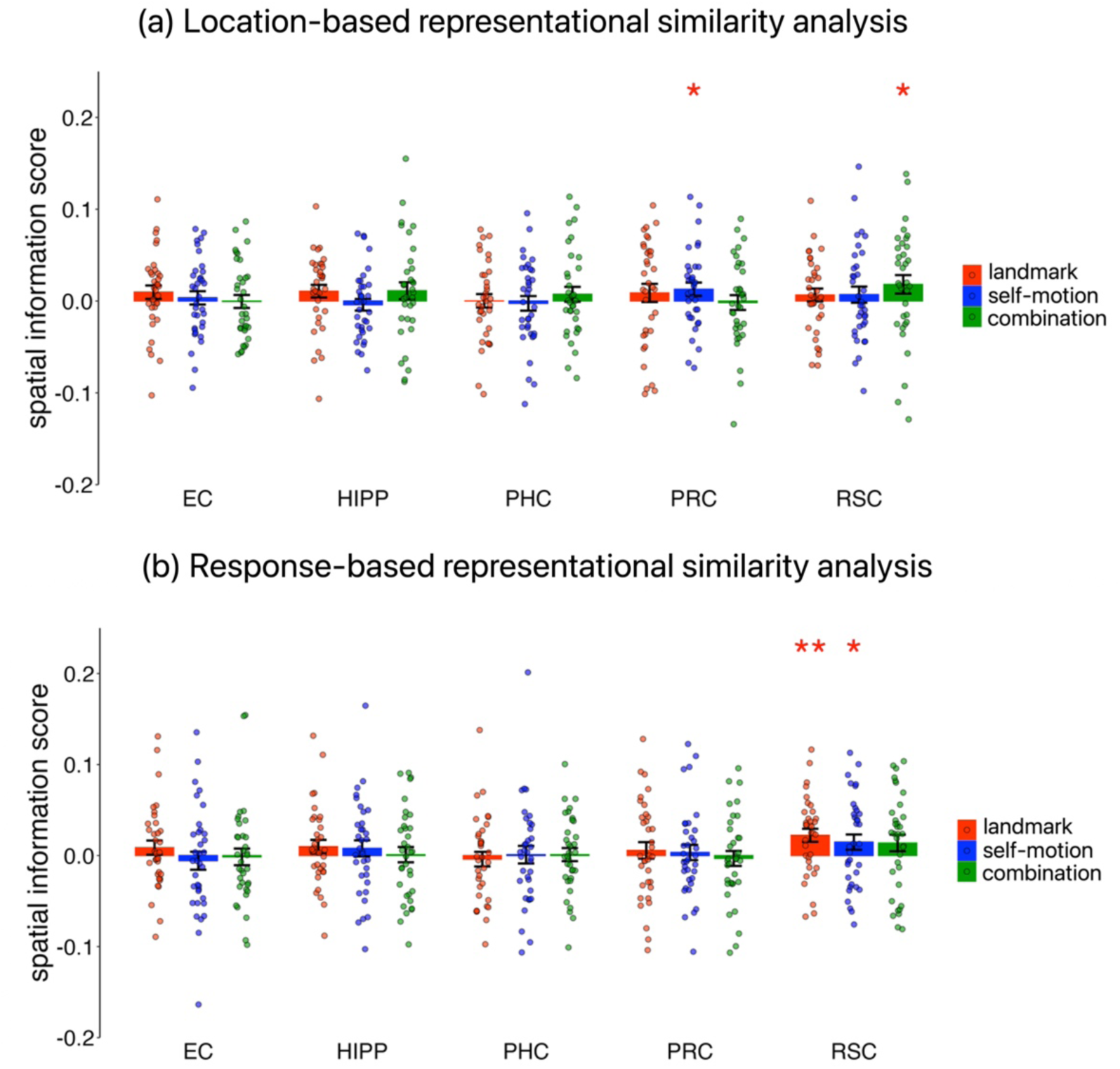
Results of representational similarity analysis. **(a)** Location-based representational analysis in all ROIs. We used representational similarity analysis to investigate whether our regions of interest contained a similar spatial coding in the form of multi-voxel pattern similarity. We hypothesized that locations spatially closer to each other would evoke more similar neural representations, as indexed by the multi-voxel activation patterns. We computed multivoxel similarities between pairs of test locations by assessing the correlations between voxel-to-voxel activation patterns. A spatial information score was derived by correlating activation pattern similarity with inter-locaiton distance (Fisher-transformed and reversed in sign). A positive spatial information score indicates that the spatial relations between test locations are encoded in the brain activity (refer to Chen et al., 2025 for detailed methodology). To preview, we did not detect any cue-integration-related coding in our ROIs.Location-based spatial information scores were tested against 0 using one-tailed simple t tests. Spatial information scores were significantly above zero in the RSC in the combination condition (t(34) = 1.7889, p = 0.041), and in PRC in the self-motion condition (t(34) = 1.751, p-value = 0.044). However, these two effects were weak and did not survive multiple comparisons correction across the 15 tests (three cue conditions * five ROIs). No other significant effects were found in any other conditions or ROIs. **(b)** Response-based representational similarity analysis. Motivated by our previous finding that RSA-based neural coding mainly reflects behavior (participant’s response) rather than stimulus (true location) (X. Chen, Wei, et al., 2025), we also analyzed response-based RSA effects. Response-based spatial information scores were significant in RSC for both the landmark (t(34) = 3.093, p_1-tailed_ = 0.002) and self-motion (t(34) = 1.752, p_1-tailed_ = 0.044) conditions. The effect in the landmark condition, but not the effect in the self-motion condition, survived multiple comparisons correction across the 15 tests (p_1-tailed,corrected_ = 0.030), indicating a preference for spatial representations derived from landmarks in RSC. No other significant effects were found in any other conditions or ROIs. Response-based spatial information score is plotted as a function of cue type (landmark vs. self-motion vs. combination) for each ROI separately (EC, HIPP, PRC, PHC and RSC). Dots represent individual participants. “*” – p_1-tailed_ < 0.05; “**” – p_1-tailed_ < 0.01. Error bars represent ±SE.

**Figure S7.**
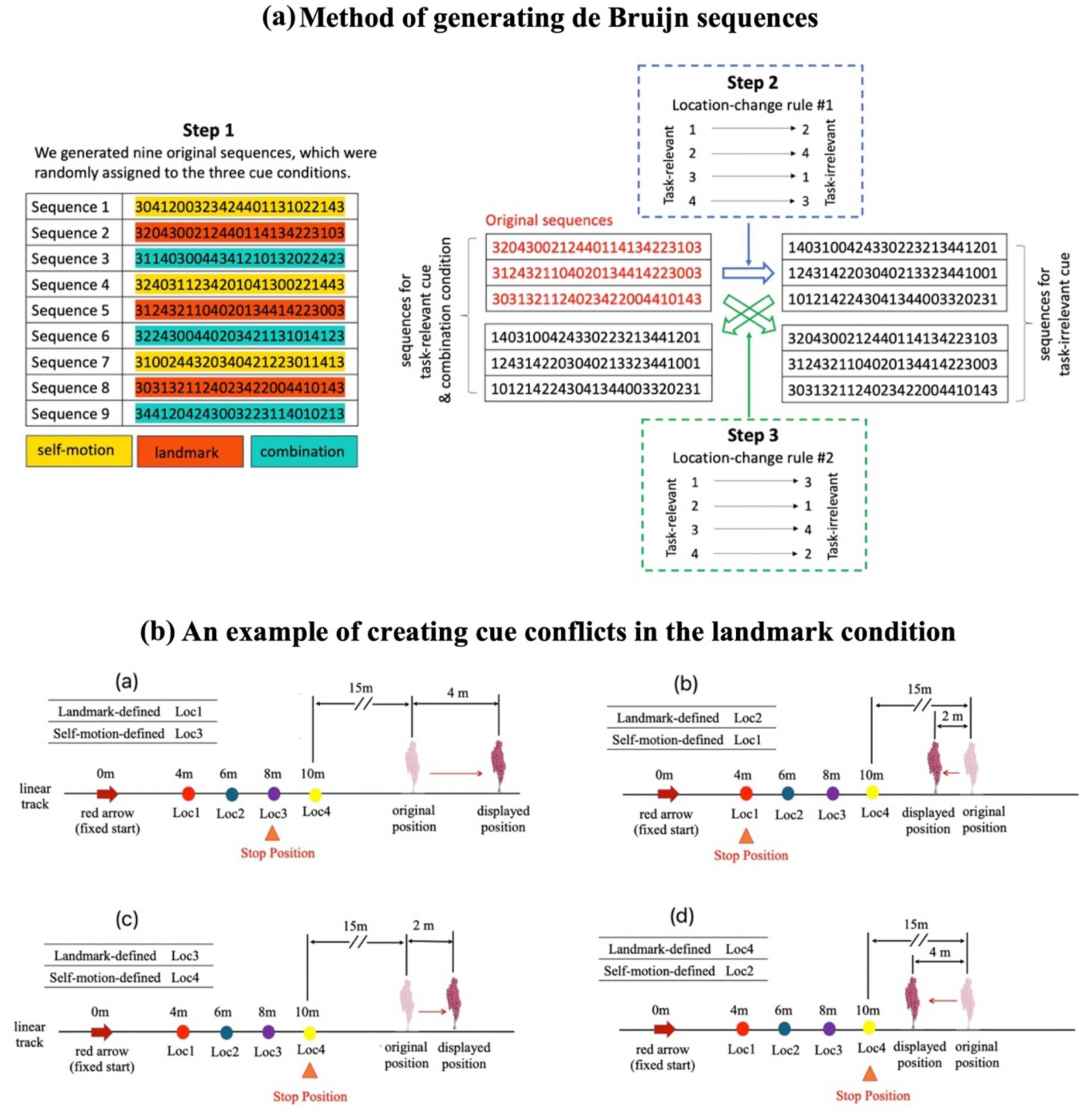
Generating de Bruijn sequences and creating cue conflicts. (a) The method of generating de Bruijn sequences. There were three cue conditions in our experiments: landmark condition, self-motion condition, and combination condition. Each condition had six blocks in the fMRI scanning. Hence, each condition needed six de Buijn sequences. Importantly, in the two single-cue conditions, the landmark cue and self-motion cues indicated different target locations. Subjects were asked to judge their locations by the landmark cue exclusively (landmark condition) or self-motion cues exclusively (self-motion condition). Because initially we were also interested in how the spatial information derived from the task-irrelevant cue type was encoded in the brain, in each single-cue block, we needed two de Brujin sequences, one for the task-relevant cue and the other for the task-irrelevant cue. We needed to make sure these two sequences had no systematic differences in detection power of adaptation. Additionally, we also needed to ensure comparable detection power of adaptation between the de Bruijn sequences utilized in different cue conditions. The procedure consisted of three steps. In Step 1, we generated nine original de Bruijn sequences with high detection power and low correlation coefficient, using the de Bruijn cycle generator. For each participant, the nine sequences were randomly assigned to the three cue conditions, with three sequences for each cue condition. In these original de Bruijn sequences, the labels 1, 2, 3, and 4 denotes the test location 1 (Loc1 at 4 vm), test location 2 (Loc2 at 6 vm), test location 3(Loc3 at 8 vm), and test location 4 (Loc4 at 10 vm). For each cue type, the three original de Bruijn sequences were used for the task-relevant cue. This step ensures comparable detection power of adaptation between the three cue conditions. In Step 2, we modified three original de Bruijn sequences to create three sequences for the concurrent task-irrelevant cue. We conceived a location-identity-change rule, such that the inter-location distance in the original sequence is least correlated with the inter-location distance in the modified sequence (Pearson r = 0.03). According to this rule, if the target location defined by the task-relevant cue is Loc 1, Loc2, Loc3, and Loc4, the corresponding target location defined by the task-irrelevant cue is Loc3, Loc1, Loc4, and Loc2. The location-identity-change rule ensured that location-based adaptation effect of the task-relevant cue and the location-based adaptation effect of the task-irrelevant cue could be dissociated. The position of the landmark was shifted according to the location-identity-change rule. In Step 3, to generate the other three de Bruijn sequences for the task-relevant cue, we used the altered de Bruijn sequences for the task-irrelevant cue from the previous step (Step 2). And to generate the other three de Buijn sequences for the task-irrelevant cue, we used the three original de Bruijn sequences for the task-relevant cue from the previous step (Step 2). This entails a new location-identity-change rule: if the target location defined by the task-relevant cue is Loc1, Loc2, Loc3, and Loc4, the corresponding target location defined by the task-irrelevant cue is Loc2, Loc4, Loc1, and Loc 3. Step 2 and Step 3 ensure comparable detection power of adaptation between the task-relevant cue and the task-irrelevant cue in the same series of trials, and at the same time, dissociability of the location-based adaptation effect between the task-relevant cue and task-irrelevant cue in the same series of trials. **(b)** An example of creating cue conflicts in the landmark condition.

**Table S1.** Results of voxel-wise analysis of location-based adaptation Displayed are significant adaptation effects for all participants, integrators only, and non-integrators only. Using the non-parametric permutation-based test and the cluster-level inference approach (cluster-defining threshold: T > 3), we conducted multiple comparisons correction across the entire search volume.

| Brain Region | MNI |  | Voxel Level<br>(T) | Cluster<br>Size (k) | pFWE-corr,<br>1-tailed |
| --- | --- | --- | --- | --- | --- |
|  | RH | LH |  |  |  |
| *All participants, combination condition* |  |  |  |  |  |
| Angular gyrus |  | -38, -60, 20 | 4.62 | 97 | 0.0084 |
|  |  | -36, -60, 32 | 4.23 |  |  |
|  |  | -46, -62, 26 | 4.21 |  |  |
| Precuneus | 6, -54, 28 |  | 4.44 | 33 | 0.0470 |
| Middle temporal gyrus |  | -58, -18, -20 | 4.22 | 69 | 0.0134 |
|  |  | -70, -22, -16 | 4.06 |  |  |
|  |  | -62, -6, -20 | 3.84 |  |  |
| *Integrators, combination condition* |  |  |  |  |  |
| Dorsal posterior cingulate | 2, -50, 30 |  | 5.73 | 81 | 0.0130 |
| Precuneus | 6, -58, 38 |  | 3.77 |  |  |
|  | 2, -58, 22 |  | 3.57 |  |  |
| Middle temporal gyrus | 64, -12, -24 |  | 4.53 | 40 | 0.0416 |
|  | 56, -12, -18 |  | 3.58 |  |  |
| Angular gyrus |  | -32, -54, 26 | 4.38 | 36 | 0.0496 |
|  |  | -38, -62, 30 | 3.89 |  |  |
| *Non-integrators, landmark condition* |  |  |  |  |  |
| Superior occipital gyrus |  | -6, -82, 44 | 4.95 | 40 | 0.0400 |
| *Non-integrators, self-motion condition* |  |  |  |  |  |
| Precuneus |  | -2, -66, 28 | 7.94 | 90 | 0.0112 |
|  |  | -10, -56, 26 | 5.14 |  |  |
|  |  | -2, -64, 38 | 4.79 |  |  |
|  |  | -30, -78, 46 | 5.47 |  |  |
| Inferior parietal lobule |  | -30, -78, 46 | 5.47 | 68 | 0.0142 |
| Angular gyrus | 48, -70, 32 |  | 4.35 | 64 | 0.0166 |
| Middle occipital gyrus | 42, -78, 38 |  | 4.32 |  |  |
|  | 48, -82, 28 |  | 3.57 |  |  |

## References

Aguirre, G. K., Mattar, M. G., & Magis-Weinberg, L. (2011). De Bruijn cycles for neural decoding. NeuroImage, 56(3), 1293–1300.

Barron, H. C., Garvert, M. M., & Behrens, T. E. (2016). Repetition suppression: A means to index neural representations using BOLD? Philosophical Transactions of the Royal Society B, 371(1705), 20150355.

Berron, D., Vieweg, P., Hochkeppler, A., Pluta, J., Ding, S.-L., Maass, A., Luther, A., Xie, L., Das, S., & Wolk, D. (2017). A protocol for manual segmentation of medial temporal lobe subregions in 7 Tesla MRI. NeuroImage: Clinical, 15, 466–482.

Bonnici, H., Chadwick, M., Kumaran, D., Hassabis, D., Weiskopf, N., & Maguire, E. A. (2012). Multi-voxel pattern analysis in human hippocampal subfields. Frontiers in Human Neuroscience, 6. https://www.frontiersin.org/journals/human-neuroscience/articles/10.3389/fnhum.2012.00290

Bromiley, P. A. (2013). *Products and convolutions of Gaussian probability density functions*. https://api.semanticscholar.org/CorpusID:18045887

Brunec, I. K., Bellana, B., Ozubko, J. D., Man, V., Robin, J., Liu, Z.-X., Grady, C., Rosenbaum, R. S., Winocur, G., Barense, M. D., & Moscovitch, M. (2018). Multiple scales of representation along the hippocampal anteroposterior axis in humans. Current Biology, 28(13), 2129–2135.e6. 10.1016/j.cub.2018.05.016

Bullmore, E. T., & Bassett, D. S. (2011). Brain graphs: Graphical models of the human brain connectome. Annual Review of Clinical Psychology, 7, 113–140. 10.1146/annurev-clinpsy-040510-143934

Burnham, K. P., & Anderson, D. R. (2002). *Model Selection and inference: A practical information-theoretic approach* (2nd Edition). Springer-Verlag. 10.1007/b97636

Campbell, M. G., Attinger, A., Ocko, S. A., Ganguli, S., & Giocomo, L. M. (2021). Distance-tuned neurons drive specialized path integration calculations in medial entorhinal cortex. Cell Reports, 36(10), 109669. 10.1016/j.celrep.2021.109669

Chadwick, M. J., Bonnici, H. M., & Maguire, E. A. (2014). CA3 size predicts the precision of memory recall. Proceedings of the National Academy of Sciences, 111(29), 10720–10725.

Chamizo, V., Manteiga, R. D., Rodrigo, T., & Mackintosh, N. J. (2006). Competition between landmarks in spatial learning: The role of proximity to the goal. Behavioural Processes, 71(1), 59–65.

Chen, G. F., King, J. A., Burgess, N., & O’Keefe, J. (2013). How vision and movement combine in the hippocampal place code. Proceedings of the National Academy of Sciences of the United States of America, 110(1), 378–383.

Chen, G., Manson, D., Cacucci, F., & Wills, T. J. (2016). Absence of visual input results in the disruption of grid cell firing in the mouse. Current Biology, 26(17), 2335–2342.

Chen, X., Chen, Y., & McNamara, T. P. (2025). Processing spatial cue conflict in navigation: Distance estimation. Cognitive Psychology, 158, 101734. 10.1016/j.cogpsych.2025.101734

Chen, X., McNamara, T. P., Kelly, J. W., & Wolbers, T. (2017). Cue combination in human spatial navigation. Cognitive Psychology, 95, 105–144.

Chen, X., Vieweg, P., & Wolbers, T. (2019). Computing distance information from landmarks and self-motion cues—Differential contributions of anterior-lateral vs. Posterior-medial entorhinal cortex in humans. NeuroImage, 202, 116074. 10.1016/j.neuroimage.2019.116074

Chen, X., Wei, Z., & Wolbers, T. (2024). Repetition suppression reveals cue-specific spatial representations for landmarks and self--motion cues in the human retrosplenial cortex. eNeuro, 11(4), ENEURO.0294-23.2024. 10.1523/ENEURO.0294-23.2024

Chen, X., Wei, Z., & Wolbers, T. (2025). Representational similarity analysis reveals cue-independent spatial representations for landmarks and self-motion cues in human retrosplenial cortex. Imaging Neuroscience, 3, imag_a_00516.10.1162/imag_a_00516

Chen, Y., & Mou, W. (2024). Path integration, rather than being suppressed, is used to update spatial views in familiar environments with constantly available landmarks. Cognition, 242, 105662. 10.1016/j.cognition.2023.105662

Chen, Z. J., He, Y., Rosa-Neto, P., Germann, J., & Evans, A. C. (2008). Revealing modular architecture of human brain structural networks by using cortical thickness from MRI. Cerebral Cortex, 18(10), 2374–2381. 10.1093/cercor/bhn003

Cheng, K., Shettleworth, S. J., Huttenlocher, J., & Rieser, J. J. (2007). Bayesian integration of spatial information. Psychological Bulletin, 133(4), 625–637.

Cisler, J. M., Bush, K., & Steele, J. S. (2014). A comparison of statistical methods for detecting context-modulated functional connectivity in fMRI. NeuroImage, 84, 1042–1052. 10.1016/j.neuroimage.2013.09.018

Cohen, N. J., & Eichenbaum, H. (1993). Memory, amnesia, and the hippocampal system. (pp. xii, 330). The MIT Press.

Cooper, R. A., & Ritchey, M. (2020). Progression from feature-specific brain activity to hippocampal binding during episodic encoding. The Journal of Neuroscience, 40(8), 1701. 10.1523/JNEUROSCI.1971-19.2019

Danon, L., Díaz-Guilera, A., & Arenas, A. (2006). The effect of size heterogeneity on community identification in complex networks. Journal of Statistical Mechanics: Theory and Experiment, 2006(11), P11010. 10.1088/1742-5468/2006/11/P11010

Diersch, N., Valdes-Herrera, J. P., Tempelmann, C., & Wolbers, T. (2021). Increased hippocampal excitability and altered learning dynamics mediate cognitive mapping deficits in human aging. The Journal of Neuroscience, 41(14), 3204–3221. 10.1523/JNEUROSCI.0528-20.2021

Doan, T. P., Lagartos-Donate, M. J., Nilssen, E. S., Ohara, S., & Witter, M. P. (2019). Convergent projections from perirhinal and postrhinal cortices suggest a multisensory nature of lateral, but not medial, entorhinal cortex. Cell Reports, 29(3), 617–627.e7. 10.1016/j.celrep.2019.09.005

Ekstrom, A. D., & Yonelinas, A. P. (2020). Precision, binding, and the hippocampus: Precisely what are we talking about? Neuropsychologia, 138, 107341. 10.1016/j.neuropsychologia.2020.107341

Epstein, R. A. (2008). Parahippocampal and retrosplenial contributions to human spatial navigation. Trends in Cognitive Sciences, 12(10), 388–396. 10.1016/j.tics.2008.07.004

Epstein, R. A., & Morgan, L. K. (2012). Neural responses to visual scenes reveals inconsistencies between fMRI adaptation and multivoxel pattern analysis. Neuropsychologia, 50(4), 530–543.

Etienne, A. S., Maurer, R., & Seguinot, V. (1996). Path integration in mammals and its interaction with visual landmarks. Journal of Experimental Biology, 199(1), 201–209.

Evensmoen, H. R., Ladstein, J., Hansen, T. I., Møller, J. A., Witter, M. P., Nadel, L., & Håberg, A. K. (2015). From details to large scale: The representation of environmental positions follows a granularity gradient along the human hippocampal and entorhinal anterior-posterior axis. Hippocampus, 25(1), 119–135. 10.1002/hipo.22357

Gallistel, C. R. (1990). The organization of learning. The MIT Press.

Garvert, M. M., Saanum, T., Schulz, E., Schuck, N. W., & Doeller, C. F. (2023). Hippocampal spatio-predictive cognitive maps adaptively guide reward generalization. Nature Neuroscience, 26(4), 615–626. 10.1038/s41593-023-01283-x

Gothard, K. M., Skaggs, W. E., & McNaughton, B. L. (1996). Dynamics of mismatch correction in the hippocampal ensemble code for space: Interaction between path integration and environmental cues. Journal of Neuroscience, 16(24), 8027–8040.

Gower, J. C., & Dijksterhuis, G. B. (2004). Procrustes problems. In Oxford Statistical Science Series. Oxford University Press.

Grady, C. L. (2020). Meta-analytic and functional connectivity evidence from functional magnetic resonance imaging for an anterior to posterior gradient of function along the hippocampal axis. Hippocampus, 30(5), 456–471. 10.1002/hipo.23164

Hooke, R., & Jeeves, T. A. (1961). “Direct search” solution of numerical and statistical problems. Journal of Association of Computing Machinery, 8, 212–229.

Jakobs, O., Langner, R., Caspers, S., Roski, C., Cieslik, E. C., Zilles, K., Laird, A. R., Fox, P. T., & Eickhoff, S. B. (2012). Across-study and within-subject functional connectivity of a right temporo-parietal junction subregion involved in stimulus–context integration. NeuroImage, 60(4), 2389–2398. 10.1016/j.neuroimage.2012.02.037

Jarsky, T., Roxin, A., Kath, W. L., & Spruston, N. (2005). Conditional dendritic spike propagation following distal synaptic activation of hippocampal CA1 pyramidal neurons. *Nature Neuroscience*, *8*(12), 1667–1676. 10.1038/nn1599

JASP Team (2023). JASP (Version 0.17.1). (n.d.). [Computer software].

Jayakumar, R. P., Madhav, M. S., Savelli, F., Blair, H. T., Cowan, N. J., & Knierim, J. J. (2019). Recalibration of path integration in hippocampal place cells. Nature, 566, 533–537.

Jeffreys H. (1961). The theory of probability (3rd ed.). Oxford University Press.

Kjelstrup, K. B., Solstad, T., Brun, V. H., Hafting, T., Leutgeb, S., Witter, M. P., Moser, E. I., & Moser, M.-B. (2008). Finite scale of spatial representation in the hippocampus. Science, 321(5885), 140–143.

Kong, X.-Z., Wang, X., Pu, Y., Huang, L., Hao, X., Zhen, Z., & Liu, J. (2017). Human navigation network: The intrinsic functional organization and behavioral relevance. Brain Structure & Function, 222(2), 749–764. 10.1007/s00429-016-1243-8

Kruskal, J., & Wish, M. (1978). *Multidimensional Scaling*. SAGE Publications, Inc. 10.4135/9781412985130

Loomis, J. M., Klatzky, R. L., Golledge, R. G., & Philbeck, J. W. (1999). Human navigation by path integration. In R. G. Golledge (Ed.), Wayfinding behavior: Cognitive mapping and other spatial processes (pp. 125–151). The Johns Hopkins University Press.

Ma, W. J., Beck, J. M., Latham, P. E., & Pouget, A. (2006). Bayesian inference with probabilistic population codes. Nature Neuroscience, 9(11), 1432–1438. 10.1038/nn1790

Madhav, M. S., Jayakumar, R. P., Li, B. Y., Lashkari, S. G., Wright, K., Savelli, F., Knierim, J. J., & Cowan, N. J. (2024). Control and recalibration of path integration in place cells using optic flow. Nature Neuroscience, 27(8), 1599–1608. 10.1038/s41593-024-01681-9

Marchette, S. A., Vass, L. K., Ryan, J., & Epstein, R. A. (2014). Anchoring the neural compass: Coding of local spatial reference frames in human medial parietal lobe. Nature Neuroscience, 17(11), 1598–1606.

McLaren, D. G., Ries, M. L., Xu, G., & Johnson, S. C. (2012). A generalized form of context-dependent psychophysiological interactions (gPPI): A comparison to standard approaches. NeuroImage, 61(4), 1277–1286.

McNamara, T. P., & Chen, X. (2022). Bayesian decision theory and navigation. Psychonomic Bulletin & Review, 29(3), 721–752. 10.3758/s13423-021-01988-9

Morgan, L. K., MacEvoy, S. P., Aguirre, G. K., & Epstein, R. A. (2011). Distances between real-world locations are represented in the human hippocampus. Journal of Neuroscience, 31(4), 1238–1245.

Nardini, M., Jones, P., Bedford, R., & Braddick, O. (2008). Development of cue integration in human navigation. Current Biology, 18(9), 689–693.

Negen, J., Bird, L.-A., & Nardini, M. (2021). An adaptive cue selection model of allocentric spatial reorientation. Journal of Experimental Psychology: Human Perception and Performance, 47(10), 1409–1429. 10.1037/xhp0000950

Newman, P. M., Qi, Y., Mou, W., & McNamara, T. P. (2023). Statistically optimal cue integration during human spatial navigation. Psychonomic Bulletin & Review, 30(5), 1621–1642. 10.3758/s13423-023-02254-w

Nichols, T. E., & Holmes, A. P. (2002). Nonparametric permutation tests for functional neuroimaging: A primer with examples. Human Brain Mapping, 15(1), 1–25.

O’Keefe, J., & Nadel, L. (1978). The hippocampus as a cognitive map. Oxford university press.

O’Reilly, J. X., Woolrich, M. W., Behrens, T. E. J., Smith, S. M., & Johansen-Berg, H. (2012). Tools of the trade: Psychophysiological interactions and functional connectivity. Social Cognitive and Affective Neuroscience, 7(5), 604–609. 10.1093/scan/nss055

Peer, M., & Epstein, R. A. (2021). The human brain uses spatial schemas to represent segmented environments. Current Biology, 31(21), 4677–4688.e8. 10.1016/j.cub.2021.08.012

Persichetti, A. S., & Dilks, D. D. (2019). Distinct representations of spatial and categorical relationships across human scene-selective cortex. Proceedings of the National Academy of Sciences, 116(42), 21312–21317. 10.1073/pnas.1903057116

Pessoa, L., Gutierrez, E., Bandettini, P. A., & Ungerleider, L. G. (2002). Neural correlates of visual working memory: fMRI amplitude predicts task performance. Neuron, 35(5), 975– 987.

Quirk, G. J., Muller, R. U., & Kubie, J. L. (1990). The firing of hippocampal place cells in the dark depends on the rat’s recent experience. The Journal of Neuroscience, 10(6), 2008– 2017.

Scheller, M., & Nardini, M. (2023). Correctly establishing evidence for cue combination via gains in sensory precision: Why the choice of comparator matters. Behavior Research Methods. 10.3758/s13428-023-02227-w

Scoville, W. B., & Milner, B. (1957). Loss of recent memory after bilateral hippocampal lesions. *Journal of Neurology*, Neurosurgery, and Psychiatry, 20(1), 11–21. 10.1136/jnnp.20.1.11

Shah, P., Bassett, D. S., Wisse, L. E. M., Detre, J. A., Stein, J. M., Yushkevich, P. A., Shinohara, R. T., Pluta, J. B., Valenciano, E., Daffner, M., Wolk, D. A., Elliott, M. A., Litt, B., Davis, K. A., & Das, S. R. (2018). Mapping the structural and functional network architecture of the medial temporal lobe using 7T MRI. Human Brain Mapping, 39(2), 851–865. 10.1002/hbm.23887

Shettleworth, S. J., & Sutton, J. E. (2005). Multiple systems for spatial learning: Dead reckoning and beacon homing in rats. Journal of Experimental Psychology: Animal Behavior Processes, 31(2), 125–141.

Squire, L. R., Stark, C. E. L., & Clark, R. E. (2004). The medial temporal lobe. In Annual Review of Neuroscience (Vol. 27, Issue Volume 27, 2004, pp. 279–306). Annual Reviews. 10.1146/annurev.neuro.27.070203.144130

Stein, B. E., & Stanford, T. R. (2008). Multisensory integration: Current issues from the perspective of the single neuron. Nature Reviews Neuroscience, 9(4), 255–266. 10.1038/nrn2331

Takahashi, H., & Magee, J. C. (2009). Pathway interactions and synaptic plasticity in the dendritic tuft regions of CA1 pyramidal neurons. Neuron, 62(1), 102–111. 10.1016/j.neuron.2009.03.007

van den Heuvel, M. P., & Sporns, O. (2013). Network hubs in the human brain. Special Issue: The Connectome, 17(12), 683–696. 10.1016/j.tics.2013.09.012

Vann, S. D., Aggleton, J. P., & Maguire, E. A. (2009). What does the retrosplenial cortex do? Nature Reviews Neuroscience, 10(11), 792–802.

Wang, J., Wang, X., Xia, M., Liao, X., Evans, A., & He, Y. (2015). GRETNA: A graph theoretical network analysis toolbox for imaging connectomics. Frontiers in Human Neuroscience, 9. https://www.frontiersin.org/journals/human-neuroscience/articles/10.3389/fnhum.2015.00386

Watanabe, M., Nakanishi, K., & Aihara, K. (2001). Solving the binding problem of the brain with bi-directional functional connectivity. Neural Networks, 14(4), 395–406. 10.1016/S0893-6080(01)00036-3

Witter, M. P., Wouterlood, F. G., Naber, P. A., & Van Haeften, T. (2000). Anatomical organization of the parahippocampal-hippocampal network. Annals of the New York Academy of Sciences, 911(1), 1–24.

Yonelinas, A. P., Ranganath, C., Ekstrom, A. D., & Wiltgen, B. J. (2019). A contextual binding theory of episodic memory: Systems consolidation reconsidered. Nature Reviews. Neuroscience, 20(6), 364–375. 10.1038/s41583-019-0150-4

Zanchi, S., Cuturi, L. F., Sandini, G., & Gori, M. (2022). Interindividual differences influence multisensory processing during spatial navigation. Journal of Experimental Psychology: Human Perception and Performance, 48(2), 174–189. 10.1037/xhp0000973

Zhang, W., & Luck, S. J. (2008). Discrete fixed-resolution representations in visual working memory. Nature, 453(7192), 233–235. 10.1038/nature06860

Zhang, W.-H., Chen, A., Rasch, M. J., & Wu, S. (2016). Decentralized multisensory information integration in neural systems. The Journal of Neuroscience : The Official Journal of the Society for Neuroscience, 36(2), 532–547. 10.1523/JNEUROSCI.0578-15.2016

Zhao, M., & Warren, W. H. (2015a). Environmental stability modulates the role of path integration in human navigation. Cognition, 142, 96–109.

Zhao, M., & Warren, W. H. (2015b). How you get there from here: Interaction of visual landmarks and path integration in human navigation. Psychological Science, 26(6), 915– 924.

